# CLSY docking to Pol IV requires a conserved domain critical for small RNA biogenesis and transposon silencing

**DOI:** 10.1101/2023.12.26.573199

**Authors:** Luisa Felgines, Bart Rymen, Laura M. Martins, Guanghui Xu, Calvin Matteoli, Christophe Himber, Ming Zhou, Josh Eis, Ceyda Coruh, Marcel Böhrer, Lauriane Kuhn, Johana Chicher, Vijaya Pandey, Philippe Hammann, James Wohlschlegel, Florent Waltz, Julie A. Law, Todd Blevins

## Abstract

Eukaryotes must balance the need for gene transcription by RNA polymerase II (Pol II) against the danger of mutations caused by transposable element (TE) proliferation. In plants, these gene expression and TE silencing activities are divided between different RNA polymerases. Specifically, RNA polymerase IV (Pol IV), which evolved from Pol II, transcribes TEs to generate small interfering RNAs (siRNAs) that guide DNA methylation and block TE transcription by Pol II. While the Pol IV complex is recruited to TEs via SNF2-like CLASSY (CLSY) proteins, how Pol IV partners with the CLSYs remains unknown. Here we identified a conserved CYC-YPMF motif that is specific to Pol IV and is positioned on the complex exterior. Furthermore, we found that this motif is essential for the co-purification of all four CLSYs with Pol IV, but that only one CLSY is present in any given Pol IV complex. These findings support a “one CLSY per Pol IV” model where the CYC-YPMF motif acts as a CLSY-docking site. Indeed, mutations in and around this motif phenocopy *pol iv* null mutants. Together, these findings provide structural and functional insights into a critical protein feature that distinguishes Pol IV from other RNA polymerases, allowing it to promote genome stability by targeting TEs for silencing.

## Introduction

Eukaryotes maintain genome segments in different chromatin states, including transcriptionally permissive euchromatin and typically silent heterochromatin. Transposable elements (TEs) are frequently marked with chromatin modifications associated with transcriptional silencing, including DNA methylation, which limits potentially dangerous transposition events and protects genome integrity^1^. In animals, this process is directed by PIWI protein-associated RNAs (piRNAs), whose precursors are synthesized by RNA polymerase II (Pol II)^2^. By contrast, plants target TEs using specialized transcription machinery, including RNA polymerase IV (Pol IV), to generate small interfering RNAs (siRNAs), and RNA polymerase V (Pol V), to produce nascent long non-coding RNAs, that mediate RNA-directed DNA methylation (RdDM)^3–5^. Notably, these two plant polymerases do not interact with the key recruitment and initiation factors of Pol II (TFIIs), but instead interact with their own polymerase-specific partners^6–12^. Thus, animal and plant lineages have converged on sequence-specific TE silencing mechanisms via distinct RNA polymerases and small RNA pathways^13–15^.

In the model plant Arabidopsis, Pol IV initiates siRNA biogenesis when recruited to chromosomal DNA via four CLASSY (CLSY) proteins (CLSY1/2/3/4)^10,16^, which have SNF2 domains similar to ATP-dependent chromatin remodelers^17,18^ and specifically interact with Pol IV^9,19^ via an unknown mechanism (**Fig. 1A**). The different CLSYs control siRNA production at distinct subsets of loci throughout the genome^10,20,21^. CLSY1 and CLSY2 function primarily in the chromosome arms, while CLSY3 and CLSY4 act primarily in pericentromeric heterochromatin^10,20^. In addition to this locus-specific targeting, the CLSYs are differentially expressed during plant development^20,22,23^ and are required for the tissue-specific regulation of DNA methylation patterns^20,23^. In all tissues tested, Pol IV targeting by CLSY1 and CLSY2 requires SAWADEE HOMEODOMAIN HOMOLOG (SHH1), a chromatin reader that binds histone tails with methylated histone 3 lysine 9 (H3K9me) and unmethylated histone 3 lysine 4 (H3K4) residues^6,19^. Mechanistically, CLSY1 and SHH1 directly interact *in vitro*^19^, and CLSY1 and CLSY2, but not CLSY3 and CLSY4, are required to mediate the interaction between SHH1 and the Pol IV complex *in vivo*^10^. Together, these findings link Pol IV targeting by CLSY1 and CLSY2 to H3K9me modifications. For CLSY3 and CLSY4, the mechanisms for locus-specific targeting remain poorly understood, but they are independent of H3K9 methylation^10^, demonstrating distinct protein interactions and modes of Pol IV targeting for CLSY1 and CLSY2 vs. CLSY3 and CLSY4. Additional accessory factors that modify Pol IV targeting and siRNA production at specific genomic sites have also been discovered, including the chromatin reader ZMP^24^, demonstrating that Pol IV recruitment is highly regulated.

**Figure 1.**
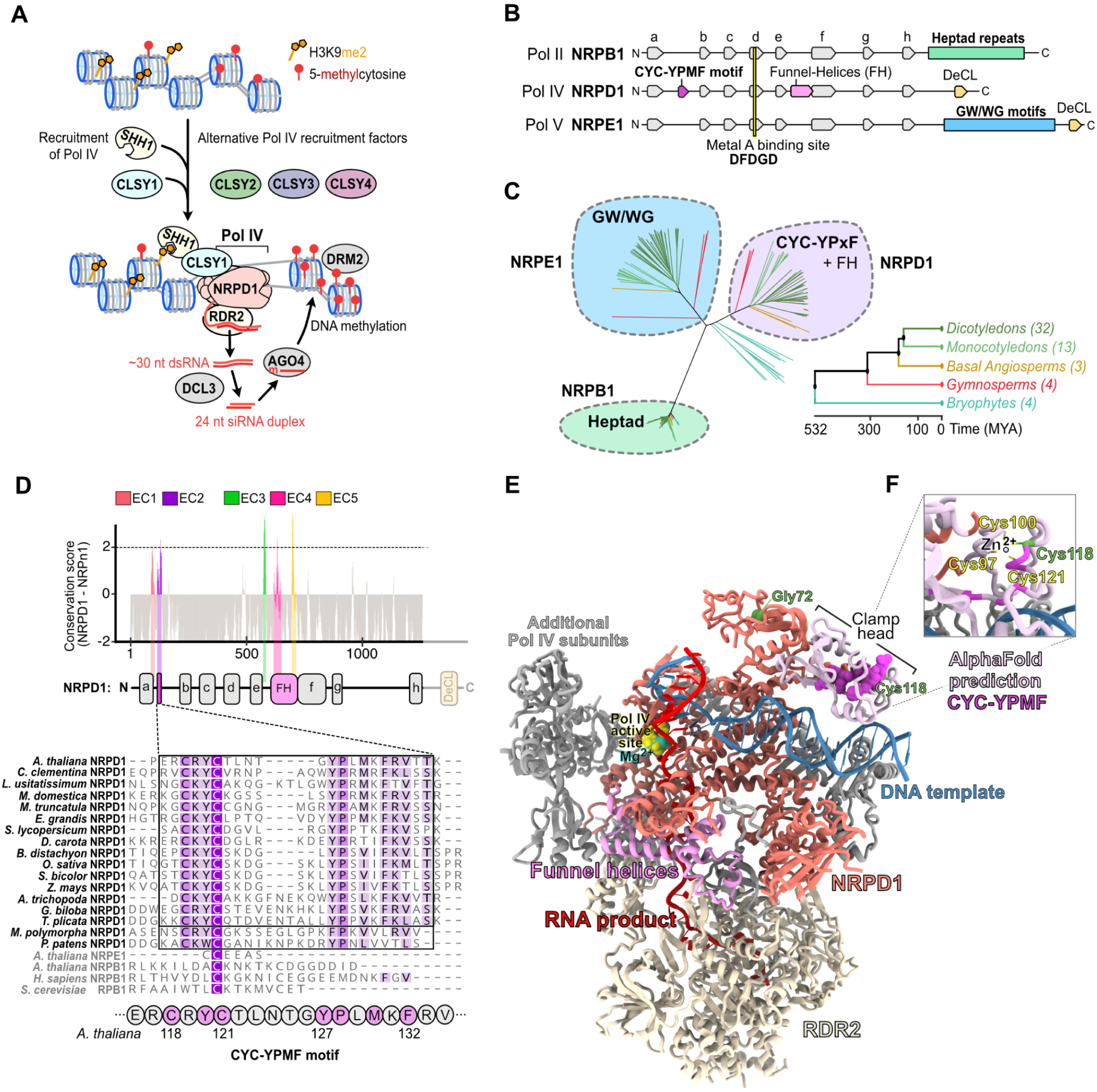
The Pol IV subunit NRPD1 has an exclusively conserved motif exposed at the enzyme exterior. **(A)** Simplified model depicting how Pol IV is recruited to loci for siRNA biogenesis and RdDM. Pol IV recruitment depends on specific partnerships with a chromatin reader SHH1 and four CLASSY proteins (CLSY1/2/3/4) in Arabidopsis. CLSY1 and CLSY2 partner with SHH1, while CLSY3 and CLSY4 function independently of SHH1. Pol IV transcripts are used by RDR2 to synthesize dsRNA that is diced by DCL3 into 24 nt siRNAs that guide DNA methylation. **(B)** Domain architectures of the largest subunits of Pol II (NRPB1), Pol IV (NRPD1) and Pol V (NRPE1). **(C)** Phylogenetic analysis of NRPB1, NRPD1 and NRPE1 subunits from 56 species with color shading indicating proteins containing heptad repeats typical of NRPB1 (green), multiple GW/WG motifs characteristic for NRPE1 (blue) and the CYC-YPxF motif typical for NRPD1 (purple). The species represent basal angiosperms, monocotyledons, dicotyledons, gymnosperms and bryophytes, with the number of species in parentheses and lines colored based on the species groupings (**Table S1**). **(D)** ConSurf analysis of NRPD1 proteins from 52 vascular plant species compared to NRPB1 and NRPE1. Five protein regions that are peaks of NRPD1-exclusive conservation (labeled EC1-EC5) are plotted in color along the subunit’s domain architecture. Positions and domains with low NRPD1 exclusive conservation are in grey. The NRPD1 C-terminal domain, including the DeCL domain, was excluded from this analysis. A multiple alignment of CYC-YPxF motifs from diverse species is shown below with conserved residues in purple. The amino acid numbering is based on Arabidopsis NRPD1. **(E)** An atomic model of the Arabidopsis Pol IV-RDR2 (7EU0) complex displayed as a cartoon representation. NRPD1 is shown mostly in coral, with the Pol IV active site in yellow, a Mg^2+^ bound at the metal A site in teal, the funnel helices in pink, the DNA template in blue and the RNA product of Pol IV in crimson red. The Pol IV clamp head, which was unmodelled in 7EU0, was predicted using AlphaFold2 and is shown in light pink, with its CYC-YPMF motif shown as an atomic sphere representation in bright purple. NRPD1 positions Gly72 and Cys118 (green) are mutated in the *nrpd1-49* and *nrpd1-50* mutants^38^, respectively. **(F)** A zoomed-in inset details the CYC-YPMF motif position (bright purple). Cys118 and Cys121 of the CYC-YPMF together with Cys97 and Cys100 of the ‘a’-domain, are predicted to coordinate a zinc ion. For better visual representation, NRPD2 is hidden. This analysis is further detailed in **Fig. S3** and **S4**.

Once targeted, Pol IV transcription is physically and enzymatically coupled to RNA-DEPENDENT RNA POLYMERASE 2 (RDR2)^9,11,25–27^. Pol IV transcribes DNA into short transcripts^11,28,29^ that serve as templates for RDR2 to synthesize a second RNA strand^25^ (**Fig. 1A**). The resulting ~30 nucleotide (nt) double-stranded RNAs (dsRNAs) are cleaved by DICER-LIKE 3 (DCL3) into 24 nt siRNA duplexes^28,29^ (**Fig. 1A**). These siRNA strands are loaded into ARGONAUTE family proteins (AGO4/6/9), which guide DNA methylation via base-pairing to nascent transcripts synthesized by another plant-specific polymerase, Pol V^30–32^.

Both Pol IV and Pol V arose from the duplication and neofunctionalization of genes that encode the 12 subunits of eukaryotic Pol II^3,33–36^. The Pol II, Pol IV, and Pol V complexes are distinguishable by their respective largest subunits, NRPB1, NRPD1, and NRPE1^4,12^. Pol IV-specific functions are thus encoded in its unique largest catalytic subunit, NRPD1 (**Fig. 1B**), and in its interactions with unique accessory components including RDR2^9,11^, SHH1^6,9^, and the CLSYs^9,10^. Indeed, cryo-electron microscopy (cryo-EM) of the Arabidopsis Pol IV complex recently revealed that RDR2 and NRPD1 are joined by NRPD1-specific funnel helices (**Fig. 1B**) that channel RNA templates into the RDR2 active site during Pol IV backtracking^26,37^. However, the features underlying other Pol IV-specific activities remain unclear. For example, it is not known how the CLSYs specifically associate with Pol IV, and it remains unclear why a point mutation in NRPD1, C118Y, that does not affect Pol IV core complex assembly or Pol IV-RDR2 association, has a global effect on siRNA levels^38^.

Taken together, previous work has demonstrated that Pol IV evolved from Pol II to couple DNA transcription to dsRNA synthesis in the context of TE-rich chromatin. However, the specific Pol IV features (e.g., amino acids, motifs, etc.) that enable these unique activities have remained elusive. To identify these features, we conducted a phylogenetic analysis of the largest subunits of RNA Pol II (NRPB1), Pol IV (NRPD1) and Pol V (NRPE1), which revealed exclusively conserved (EC) regions in NRPD1. Of these five EC regions, three (EC3-5) mapped at or near the funnel helices that connect NRPD1 to RDR2 and allow the rapid conversion of Pol IV transcripts into dsRNAs^25,26^. This validates our approach for finding NRPD1-specific features linked to unique Pol IV activities. By leveraging structural, biochemical, and molecular assays we were then able to map the EC1 and EC2 regions and reveal a key role for EC2 in facilitating Pol IV targeting. Specifically, we found that EC1 and EC2 occupy the previously unresolved Pol IV clamp head region, where four cysteines come together to coordinate a zinc ion: two from EC1 and two from the CYC-YPMF motif^38^ identified here as part of EC2. Via a series of immunoprecipitation and mass spectrometry experiments we found that the CYC-YPMF motif is required for the interaction between Pol IV and the CLSYs and that only one CLSY family member is present in any given Pol IV complex. Together, these findings supporting a model in which the CYC-YPMF motif in EC2 acts as a CLSY-docking site. Consistent with these findings, mutations in the CYC-YMPF motif, or in another conserved residue in the neighboring clamp core region, resulted in global losses of siRNAs and defects in TE silencing. Together, these structure-function studies identified an exclusively conserved domain within NRPD1 that distinguishes Pol IV from Pol II and Pol V and acts as a CLSY docking site that allows Pol IV to target, transcribe and silence TEs throughout the genome.

## Results

### Identification of five exclusively conserved regions in the largest subunit of Pol IV

To detect protein motifs that represent unique features of Pol IV, we compared the largest subunits of Pol II, Pol IV and Pol V from diverse plant species. BLASTP queries of the Arabidopsis proteins AtNRPB1, AtNRPD1 and AtNRPE1 against the Phytozome13 and NCBI databases identified numerous close homologs. To pass our quality criteria, we required these proteins to be over 900 amino acids long and contain the catalytic metal A binding site residues (DFDGD) common to all RNA polymerase largest subunits^39–41^, resulting in a total of 202 protein sequences from 56 species (**Fig. 1C**, **S1 and S2A**; **Table S1**). Phylogenetic analysis based on MUSCLE alignment grouped these subunits of Pol II, Pol IV and Pol V (NRPB1/D1/E1) into four clusters. One clade of 74 protein sequences clustered around AtNRPB1, all of which had heptad repeats in their C-terminal domain (CTD), indicating that they are the largest subunits of Pol II (**Fig. 1C**, green). As expected given the divergence in the transcription cycles of Pol IV and Pol V compared to Pol II^5,31,42^, none of the NRPD1 or NRPE1 proteins have the heptad repeats associated with NRPB1 (**Fig. 1C, Table S1**). Instead, these polymerase subunits are part of two independent clades. AtNRPD1 is in a clade including 58 proteins which contain CYC-YPMF like motifs (CYC-YPxF) of unknown function^38^, as well as the amino acids corresponding to funnel helices that connect the catalytic domains of Pol IV and RDR2 in Arabidopsis^26^ (**Fig. 1C**, purple background). AtNRPE1 is in another clade encompassing 58 sequences that contain repeated GW/WG motifs in their CTDs (**Fig. 1C**, blue). In Arabidopsis, Pol V’s GW/WG motifs facilitate the interaction between Pol V and siRNA-loaded AGO proteins to guide *de novo* DNA methylation^30,43,44^.

In contrast to polymerases from the abovementioned vascular plants, the bryophyte NRPD1/NRPE1-like sequences form a distinct, fourth cluster of 12 proteins in our phylogenetic tree (**Fig. 1C**; teal branches, no background color). One group of proteins in this cluster contains the largest subunit of *Physcomitrium patens* Pol IV (**Fig. S2A**, Ppa_NRPD1)^45^. In three bryophyte species, more than one NRPE1-like protein was detected (**Fig. S2A**). The bryophyte NRPE1-like proteins typically have a shortened CYC-YP motif with divergent amino acids between their ‘a’ and ‘b’-domains, in addition to the expected CTD GW/WG motifs (**Fig. S2A**). However, none of the bryophyte NRPD1 or NRPE1-like proteins had a fully intact CYC-YPxF motif (**Fig. S2A**). In light of the solid molecular evidence for AtNRPD1’s specific role in Arabidopsis Pol IV, ZmNRPD1’s specific role in maize Pol IV and OsNRPD1’s specific role in rice Pol IV^12,46–48^, our phylogenetic analyses suggest that the NRPD1 CYC-YPxF motif arose as a functionally important Pol IV amino acid sequence in a common ancestor of the vascular plants but after their split from the bryophytes.

Having clustered the largest subunits of all three RNA polymerases, we next sought to identify regions that are specifically conserved in NRPD1, as compared to NRPB1 or NRPE1. To this end, we measured the evolutionary conservation^49^ of each AtNRPD1 amino acid position relative to corresponding NRPB1 and NRPE1 positions across all the orthologs identified from vascular plants. This analysis identified five major peaks of Pol IV-exclusive conservation (EC1-5), where most amino acids in a nine amino acid window are conserved in NRPD1 but not in NRPB1 or NRPE1 (**Fig. 1D**). One of these regions, EC1, partly overlaps domain ‘a’, the first of eight domains that are broadly conserved in the largest subunits of Pol II, Pol IV and Pol V (**Fig. 1D**, lowercase ‘a’ to ‘h’)^5,50^, while the others are in regions conserved only in Pol IV (**Fig. 1D** and **S2B**). Three of the EC regions (EC3, EC4 and EC5) are between the NRPD1 ‘e’ and ‘f’-domains (**Fig. 1D**). While EC3 is a previously unexamined Pol IV-conserved region, EC4 and EC5 overlap the funnel helices that contribute to RDR2’s specific association with Pol IV, rather than with Pol II or Pol V^9,11,25,26^. The other two EC regions neighbor each other in the NRPD1 N-terminus, one overlapping part of the NRPD1 ‘a’-domain (EC1) and the other covering the CYC-YPxF motif (EC2). While the mechanistic roles of EC1, EC2, and EC3 remain unknown, their strong conservation and prior data showing global reductions in siRNA levels caused by a cysteine to tyrosine mutation in EC2 (YYC-YPMF)^38^, suggest that they could impart functions that distinguish Pol IV from Pol II and Pol V.

### The CYC-YPMF motif is part of the NRPD1 clamp head on the exterior of Pol IV

To visualize the three-dimensional (3D) locations of the five EC domains within AtNRPD1, we sought to map them onto the cryo-EM structure of the Arabidopsis Pol IV-RDR2 complex^26^ (7EU0). Overall, the AtNRPD1 tertiary structure in 7EU0 is similar to the largest subunits of yeast or mammalian Pol II (RPB1)^26,51–53^. However, the portion of NRPD1 spanning EC1 and EC2, including the CYC-YPMF motif, is unresolved in the 7EU0 structure of the Pol IV-RDR2 complex. Thus, we used the AlphaFold2 prediction of NRPD1 (AF-Q9LQ02-F1) to situate this portion of NRPD1 within the Pol IV-RDR2 complex using the 7EU0 cryo-EM density data (**Fig. 1E** and **S3**). Several lines of evidence support the validity of this positioning. First, the highly conserved ‘a’ and ‘b’-domains that are directly adjacent to the CYC-YPMF motif in AtNRPD1, and form the Pol IV ‘clamp core’ domain (**Fig. S3A**, green/turquoise residues), are positioned comparably in the 7EU0 structure to those in the AlphaFold2 model of AtNRPD1 (**Fig. S3A**, orange/pink residues). Second, the confidence scores from AlphaFold2 at the CYC-YPMF residues are high (**Fig. S3B**, pLDDT) and the fit of this predicted domain into the experimental Coulomb potential map of Pol IV supports the presence of this globular domain protruding from the clamp core (EMD-31305; **Fig. S3C**). Third, the similarly positioned ‘clamp head’ domain in Pol II (human 6GMH, yeast 7O75)^26,53^ further reinforces our attribution of this density to the Pol IV region containing the CYC-YPMF. Finally, the electrostatic potential of the putative Pol IV clamp head (**Fig. 1E** and **S3A**, purple/pink residues) is positively charged on the surface facing the downstream DNA template, as expected for a domain interacting with DNA (**Fig. S3D**).

Modeling the region of NRPD1 containing EC1 and EC2 within the Pol IV-RDR2 structure (**Fig. 1E**) allowed visualization of all five EC peaks (**Fig. 1D**) on the 3D structure of NRPD1, either as a heatmap based on conservation level (**Fig. S4A**) or as discretely colored regions (**Fig. S4B**). Consistent with their proximity to the NRPD1 funnel helix region, the EC3, EC4 and EC5 regions fold into loops or helices that funnel Pol IV RNA transcripts to RDR2. The EC3 loops are adjacent to the primary RNA transcript as it exits Pol IV and enters RDR2, whereas EC4 and EC5 contain highly conserved NRPD1 amino acids that directly contact RDR2 (N638, E642, Y645, D710, L714)^26^ (see **Fig. S2B**), which validates our method for detecting Pol IV-specific features (**Fig. S4C**, bottom inset). Interestingly, we found that two universally conserved amino acids in the RNA polymerase ‘a’-domain, Cys97 and Cys100, which are imbedded in EC1, combine with Cys118 and Cys121 from EC2 to form the Pol IV clamp head (**Fig. 1E** and **S4D**, top inset). Supporting this arrangement, Cys97-Cys100-Cys118-Cys121 likely coordinate a zinc ion together, as detected using Zincbindpredict and AlphaFill (**Fig. 1F**, zoomed-in inset)^54,55^. The deep conservation of Pol IV-specific Cys118 and Cys121, along with the adjacent YPMF residues, strongly indicates this domain’s functional importance to Pol IV. Indeed, an individual C118Y mutation (*nrpd1-50*) is sufficient to disrupt 24 nt siRNA biogenesis and RdDM^38^. Despite its importance, the function of this motif was not determined^38^. Given that the CYC-YPMF motif is located in the clamp head (**Fig. 1E** and **S3A**, purple/light pink domain), which is distant from both the Pol IV active site (**Fig. 1E**, yellow aspartic acid triad) and the NRPD1 funnel helices that couple Pol IV to RDR2 (**Fig. 1E**, central pink residues), it is unlikely to play a direct role in mediating the catalytic activities of Pol IV. Instead, as the CYC-YPMF motif is exposed on the enzyme exterior, we hypothesize that it could facilitate interactions with Pol IV-specific factors that are important for the enzyme’s role in RdDM.

### Pol IV association with SHH1 and CLSY recruitment factors requires CYC-YPMF

To test whether the CYC-YPMF motif mediates Pol IV assembly with partner proteins, we transformed *nrpd1-3* mutants with constructs encoding one of three variants of the NRPD1 subunit tagged with a 3xFLAG (3xF) epitope: a wild-type variant (NRPD1-3xF_WT_), a variant with a CYC to AAA mutation (NRPD1-3xF_AAA-YPMF_) and a variant with a YPMF to AAAA mutation (NRPD1-3xF_CYC-AAAA_) (**Fig. 2A**). For each variant, three independent lines expressing similar levels of NRPD1-3xF were selected (**Fig. S5A**) and flower tissue was collected for immunopurification and mass spectrometry (IP-MS) experiments (**Fig. S5B** and **Fig. 2B, C, D**). In total, six IP-MS datasets were obtained for each NRPD1-3xF variant (i.e., two replicates for each of the three independent lines (**Table S2**)). For all six experiments, the protein spectral counts for each individual component of the purified Pol IV complexes were displayed using balloon plots and colored based on the significance of their enrichment compared to the NRPD1-3xF_WT_ control (**Fig. 2B**). These plots demonstrate that RDR2, as well as the great majority of Pol IV core subunits (NRPD1 to NRPD12), were observed in all NRPD1-3xF IP-MS replicates (WT, AAA-YPMF or CYC-AAAA; the only subunits not detectable in all replicates were some of the smaller subunits, which could be explained by their digestion into a smaller amount of observable peptides (**Fig. 2B**)). When the replicate experiments were visualized in aggregate using volcano plots, the Pol IV subunits (green dots) and RDR2 (yellow dot) clustered near the base of the volcano plot (**Fig. 2C, D**), demonstrating that the Pol IV-RDR2 complex assembles even when the CYC-YPMF motif is mutated. Indeed, of these proteins, the only one passing both the fold change (|log_2_FC| ≥ 1) and p-value (≤ 0.05) cutoffs was NRPD7b, which is enriched, rather than depleted, in the NRPD1-3xF_CYC-AAAA_ IP-MS experiments (**Fig. 2B, D**). By contrast, all four CLSY proteins were significantly depleted, and SHH1 was also depleted, though not quite passing the adj. p-value cutoff, in the NRPD1-3xF_CYC-AAAA_ and NRPD1-3xF_AAA-YPMF_ IP-MS experiments compared to the NRPD1-3xF_WT_ control (**Fig. 2B-D**). Indeed, no CLSY or SHH1 peptides were identified in IP-MS experiments using the NRPD1-3xF_CYC-AAAA_ or NRPD1-3xF_AAA-YPMF_ variants. Taken together, these data suggest that both the CYC and YPMF sub-motifs of the clamp head are required for Pol IV association with the four CLSY proteins (CLSY1, CLSY2, CLSY3 and CLSY4), as well as the chromatin reader SHH1.

**Figure 2.**
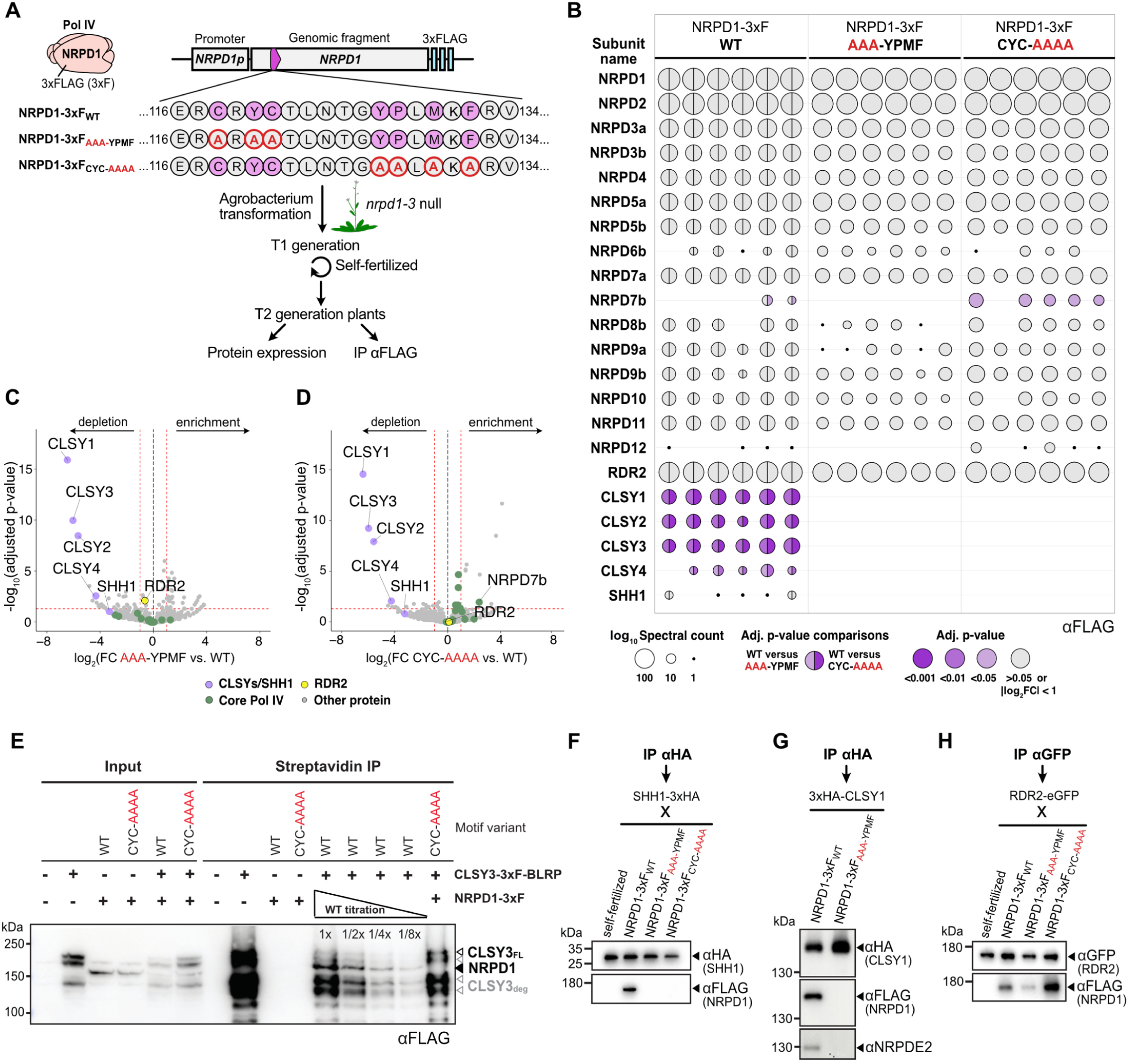
The CYC-YPMF motif is essential for copurification of Pol IV with its recruitment factors. **(A)** Experimental setup to evaluate the effect of mutations in the CYC-YPMF motif on the co-immunoprecipitation of proteins associated with the Pol IV complex (pink) using the NRPD1-3xF variants indicated below. The wild-type AtNRPD1 CYC-YPMF amino acids are indicated in purple and the amino acids altered in the other variant lines are outlined in red. These constructs were transformed into the *nrpd1-3* null mutant and experiments were conducted in the T_2_ generation. **(B)** Balloon plots showing the results from co-immunoprecipitation and mass spectrometry (IP-MS) experiments using flower extracts from three independent lines, each with two technical replicates, expressing the NRPD1-3xF_WT_, NRPD1-3xF_AAA-YPMF,_ or NRPD1-3xF_CYC-AAAA_ variants. The balloon plot shows the peptide spectral counts for Pol IV-RDR2 subunits (NRPD1 to NRPD12, RDR2) and other interactors (CLSY1-4 and SHH1) in for each dataset. The spectral count is indicated by the log_10_-transformed balloon area. Colored balloons (purple shades) represent subunits where comparisons between the NRPD1-3xF_WT_ data and either the NRPD1-3xF_AAA-YPMF_ or the NRPD1-3xF_CYC-AAAA_ variant data pass fold change (FC) and adjusted (adj.) p-value cut-offs, indicating their significance. By contrast, comparisons with |log_2_FC| < 1 or adj. p-value > 0.05 are all shown in grey. In the NRPD1-3xF_WT_ data, the left half-balloon colors are adj. p-values for comparison to the NRPD1-3xF_AAA-YPMF_ IP-MS data, and the right half-balloon colors are adj. p-values for comparison to the NRPD1-3xF_CYC-AAA_ IP-MS data. **(C and D)** Volcano plots showing the enrichment or depletion of proteins from six independent IP-MS experiments comparing the NRPD1-3xF_WT_ lines with either the NRPD1-3xF_AAA-YPMF_ or the NRPD1-3xF_CYC-AAA_ lines. The red hashed lines demarcate a log_2_FC of magnitude 1 and an adj. p-value of 0.05. Purple dots indicate Pol IV recruitment factors (CLSY1-4 and SHH1), green dots the Pol IV core subunits, and a yellow dot represents RDR2. **(E)** Anti-FLAG western blot detecting NRPD1 and CLSY3 proteins using the genotypes indicated above each lane from either input or streptavidin IP samples. Unless marked, all samples were loaded at 1x. The bands corresponding to NRPD1 as well as both full length CLSY3 (CLSY3_FL_; Black) and several CLSY3 degradation products (CLSY3_deg_; Grey) are indicated on the right. **(F)** Western blots detecting SHH1 and NRPD1 proteins in an Anti-HA IP using flowers from F1 plants of the self-fertilized SHH1-3xHA line, or flowers from this line crossed to the NRPD1-3xF variants indicated. **(G)** Western blots detecting CLSY1, NRPD1 and NRPD2 in an Anti-HA IP using flowers from 3xHA-CLSY1 plants supertransformed with the NRPD1-3xF variants indicated. **(H)** Western blots detecting RDR2 and NRPD1 proteins in an Anti-GFP IP using flowers from F1 plants of the self-fertilized RDR2-eGFP line, or flowers from this line crossed to the NRPD1-3xF variants indicated. In panels **E**-**H,** protein sizes are indicated in kDa on the left of the western analyses.

To confirm the importance of the NRPD1 CYC-YPMF motif in mediating Pol IV’s interactions with its recruitment factors via an orthogonal method, smaller scale co-immunoprecipitation (co-IP) experiments were also conducted. For these experiments, transgenes expressing CLSY3-3xF-BLRP^20^, 3xHA-CLSY1, SHH1-3xHA, or RDR2-GFP were introduced into plant lines expressing the NRPD1-3xF variants (WT, CYC-AAAA, or AAA-YPMF) via genetic crossing or super-transformation. For CLSY3, the interaction with NRPD1-3xF_WT_ but not with NRPD1-3xF_CYC-AAAA_ was confirmed in several replicate experiments (**Fig. 2E** and **S5C**). Moreover, a dilution series of the NRPD1-3xF_WT_ co-IP demonstrated a greater than 8x reduction in the interaction between CLSY3-3xF-BLRP and NRPD1-3xF_CYC-AAAA_ (**Fig 2E**). Likewise, the 3xHA-tagged SHH1 and CLSY1 proteins were each found to co-IP with NRPD1-3xF_WT_ but not with the motif mutant versions of NRPD1-3xF (**Fig. 2F, G**). These findings, along with the ability of RDR2-GFP to associate with both the WT and mutant NRPD1 variants (**Fig. 2H**), are consistent with the IP-MS results and further show the critical role of the CYC-YPMF motif in mediating interactions between Pol IV and its targeting factors—the CLSYs and SHH1.

### Pol IV complexes are demarcated by distinct CLSY proteins

Despite recent advances in our understanding of the structure and function of the core Pol IV complex, it remains unknown whether multiple different CLSYs associate with a given Pol IV complex or if each complex contains a dedicated CLSY protein. To address this question, IP-MS experiments were conducted using flower tissue collected from transgenic lines expressing 3xF-tagged versions of all four CLSYs driven by their endogenous promoters and introduced into their respective *clsy* mutant backgrounds to verify functional complementation^20^ (**Fig. 3A, B, Fig. S6** and **Table S3**). Although multiple attempts were made using both N- and C-terminally tagged CLSY2 lines, neither CLSY2 nor Pol IV subunits were detected in IP-MS experiments, suggesting either the abundance of CLSY2 is too low or the epitope is not accessible. However, purifications of the other three CLSYs were robust, as peptides from RDR2 and many Pol IV subunits were significantly enriched across several biological and technical replicates compared to non-transgenic wild-type controls (**Fig. 3A, B**). Consistent with previous work showing interactions between CLSY1 and SHH1^6,9,10,19^, peptides from SHH1 were enriched only in the CLSY1 IP-MS experiments (**Fig. 3**). Given the high quality and specificity of these CLSY IP-MS experiments, the failure to detect peptides for more than one CLSY (**Fig. 3A, B**) supports a “one CLSY per Pol IV complex” model.

**Figure 3.**
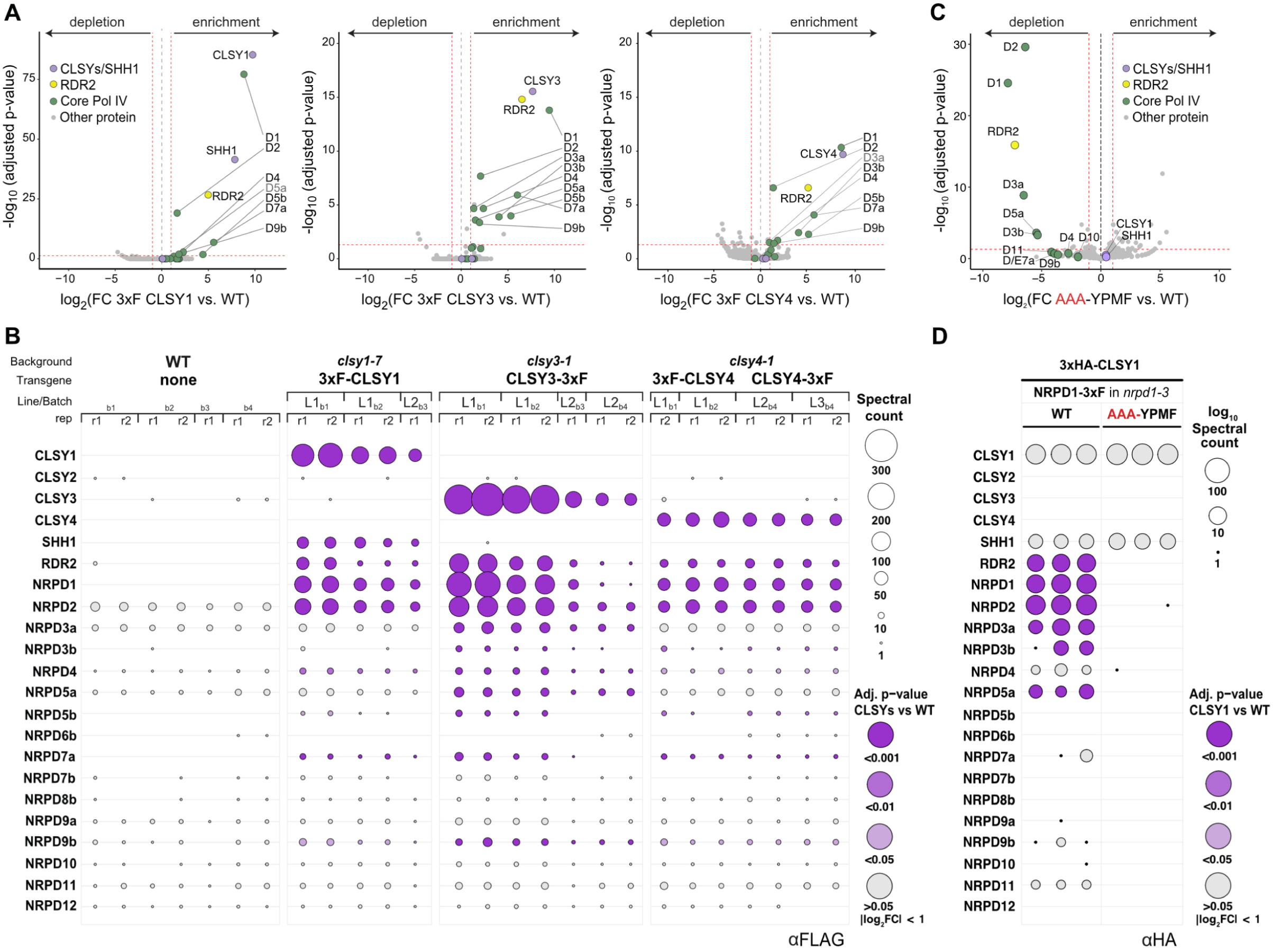
CLSY1 bridges SHH1 to Pol IV through the CYC-YPMF motif. **(A)** Volcano plots showing the enrichment or depletion of proteins from a combination of several independent CLSY1, CLSY3, and CLSY4 IP-MS experiments compared to non-transgenic control IP-MS experiments. The red hashed lines demarcate a log_2_FC of magnitude 1 and an adj. p-value of 0.05. Enriched Pol IV subunits are designated by subunit number (D1, D2, D3, …, D12) and are colored in green while SHH1 and the CLSYs are colored in light purple and RDR2 in yellow. **(B)** Balloon plot representation of the individual CLSY IP-MS experiments comparing copurifying peptides with their WT controls. The spectral counts are indicated by the linear balloon area, and significantly enriched subunits (log_2_ FC >1) for the WT vs. 3xF tagged CLSY1, CLSY3, or CLSY4 comparisons are colored based on their adj. p-values (grey to purple shades as detailed in Fig. 2B). Above each plot, the genetic background and transgenes used for each IP are indicated. The line numbers (L#) represent independent tagged lines, while the batches (b#) represent independent IP-MS experiments, most of which also included technical replicates (r#). (**C**) Volcano plot as described in (**A**) showing the enrichment or depletion of proteins from 3xHA-CLSY1 IP-MS experiments using flowers from the *nrpd1-3* mutant line complemented with either the NRPD1-3xF_WT_ or the NRPD1-3xF_AAA-YPMF_ variant. **(D)** Balloon plot representation of 3xHA-CLSY1 IP-MS experiments comparing copurifying peptides of 3xHA-CLSY1 in NRPD1-3xF/*nrpd1-3* lines using either the NRPD1-3xF_WT_or NRPD1-3xF_AAA-YPMF_ variants. The spectral counts are indicated by the log-transformed balloon area, and adj. p-values for the comparisons between the NRPD1-3xF_WT_ vs. NRPD1-3xF_AAA-YPMF_IP-MS experiments are indicated by the balloon color (grey to purple shades as detailed in Fig. 2B).

The aforementioned model, as well as the importance of the CYC-YPMF motif in mediating interactions between Pol IV and its targeting factors, is further supported by an independent set of IP-MS experiments. These experiments utilized flower tissue collected from *nrpd1-3* mutant lines that express either the NRPD1-3xF_WT_ or the NRPD1-3xF_AAA-YPMF_ variant and were super-transformed with constructs encoding 3xHA-tagged CLSY1 proteins (**Fig. 3C, D** and **Table S4**). 3xHA-CLSY1 and SHH1 peptides were recovered in anti-HA IPs in the presence of either NRPD1-3xF variant (**Fig. 3C, D**), but no other CLSYs were immunoprecipitated (**Fig. 3D**), supporting the one CLSY per complex model. Furthermore, the lack of peptides for Pol IV subunits and RDR2 specifically in the IP-MS experiments using the NRPD1-3xF_AAA-YPMF_ variant (**Fig. 3 C, D**) reinforces the importance of the CYC-YPMF motif in mediating the interaction between CLSY1-SHH1 and Pol IV. Taken together, these findings and those presented in **Fig. 2**, reveal the mechanism that prevents more than one CLSY from associating with Pol IV simultaneously: they all compete for a common NRPD1 docking site comprised of the CYC-YPMF motif. We have thus designated this motif as the “CLSY-docking” motif.

### The CYC-YPMF motif is essential for siRNA production, DNA methylation and TE silencing

To determine the impact of the CYC and YPMF mutations on the activity of the RdDM pathway, siRNA levels were assessed. Specifically, we sought to determine how the severity of these CYC-YPMF mutants compared to each other and to previously identified point mutations at or near this motif within NRPD1^38^. Furthermore, we wondered whether any of these mutants would display selective effects on siRNA clusters controlled by specific CLSY family members. To address these questions, small RNA sequencing (smRNA-seq) experiments were conducted using flower tissue from the three NRPD1-3xF variant lines (WT, CYC-AAAA, or AAA-YPMF) as well as WT, *nrpd1-3*_null_, and *clsy* quadruple (*clsy quad*) mutant controls. These data sets were then compared to a reanalysis of tissue-matched smRNA-seq experiments from two previously identified NRPD1 point mutants (*nrpd1-49*_G72E_ in the clamp core and *nrpd1-50*_YYC-YPMF_ in EC2, the clamp head), along with their respective WT and *nrpd1-51_null_*controls^38^ (**Table S5**). For the aforementioned comparisons, we conducted our analyses using the previously identified 12,939 siRNA clusters, which were categorized based on their *clsy*-dependencies^20^. To confirm the validity of using these clusters, we first demonstrated that they contain >92% of all the 24 nt siRNAs from the control samples in the two datasets being compared (92.9% for SucSul_WT_ and 92.7% for Col-0_WT_; **Fig. S7A**). Furthermore, we demonstrated that ~99% of these clusters were significantly downregulated (log_2_FC ≤ 1 and FDR < 0.01) in the *clsy* quad and both *nrpd1* null mutants, confirming that these clusters are both CLSY and Pol IV-dependent in the experiments being compared (**Fig. S7B**).

We then determined the relative strengths of the NRPD1-3xF variants and *nrpd1* point mutations across the full set of 12,939 loci via smRNA-seq experiments (**Fig. 4A** and **S7B**), or at select loci via northern blotting (**Fig. S7C** and Ferrafiat et al. 2019^38^), again relying on the vetted siRNA clusters (**Fig. S7D**). While all the mutants showed strong reductions in siRNAs, nearing the severity of *nrpd1* null mutants, some were stronger than others: the NRPD1-3xF_CYC-AAAA_ variant was the strongest, followed closely by the *nrpd1-49*_G72E_ and NRPD1-3xF_AAA-YPMF_ mutants, with the *nrpd1-50*_YYC-YPMF_ mutant being the weakest (**Fig. 4A** and **Fig. S7B, E**). These data show that the CLSY-docking motif is required for the biogenesis of nearly all Pol IV-dependent siRNAs, with the location of the mutation affecting the severity of the siRNA defects.

**Figure 4.**
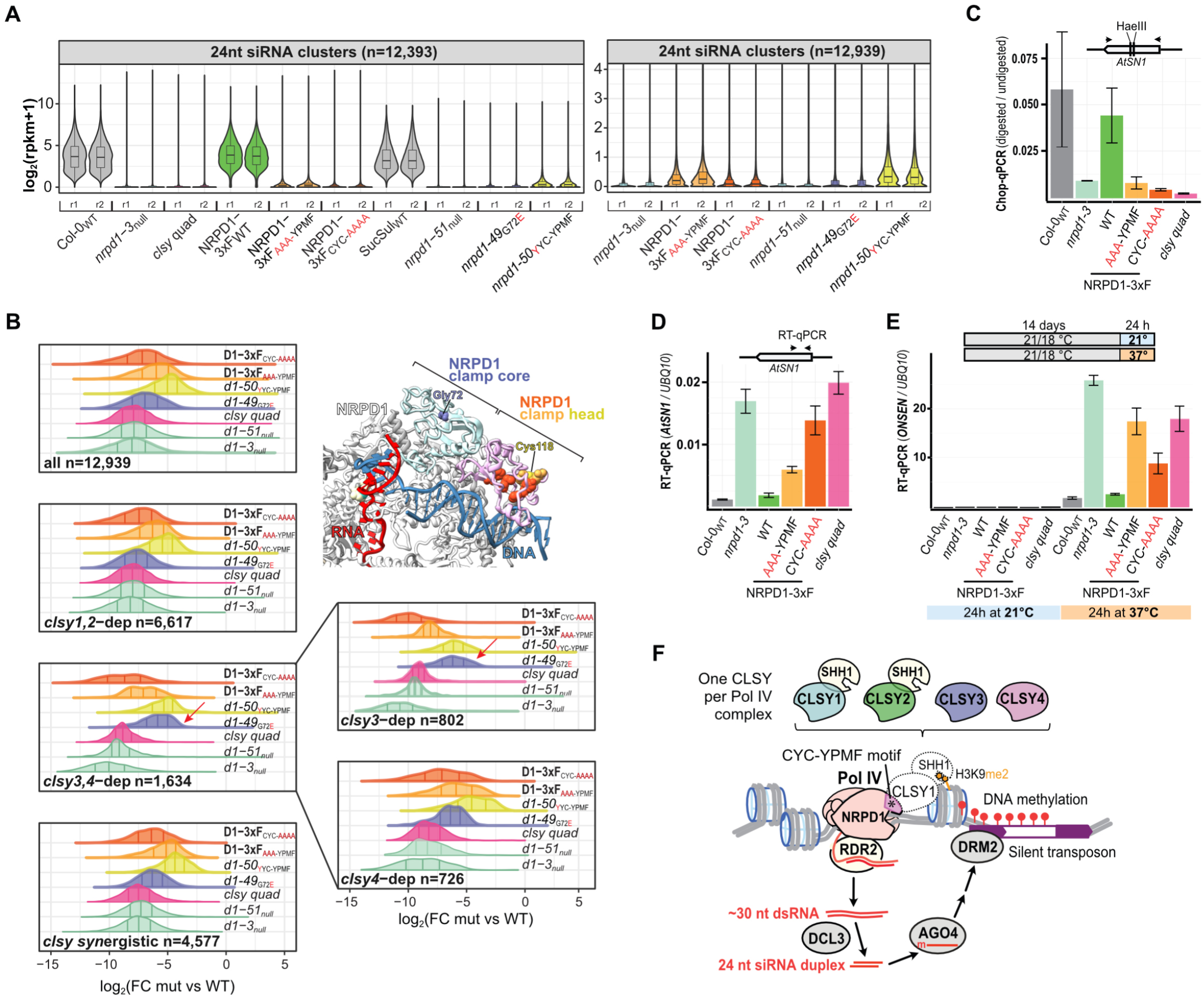
The CLSY-NRPD1 interaction is important for siRNA biogenesis and TE silencing. (**A**) Violin plots showing the normalized expression levels [log_2_(rpkm+1)] of 24 nt siRNAs across the full set of clusters (n=12,939) in the genotypes indicated below for two biological replicates (r1 and r2). Select genotypes are shown on a smaller Y-axis scale on the right. (**B**) Half violin plots showing the differential expression profiles (log_2_FC) of siRNA clusters in the indicated categories (lower left) for each mutant (right) relative to their respective wild-type controls. The NRPD1 and *nrpd1* symbols are abbreviated as D1 and *d1*, respectively. Red arrows highlight the siRNA expression profiles that are shifted relative to the rest of the categories for the *nrpd1-49*_G72E_ mutant. A portion of the Pol IV structure showing the locations of the characterized mutants is shown as an inlay. (**C**) Methylation at the *AtSN1* retroelement evaluated by Chop-qPCR. Genomic DNA was digested with a methylation-sensitive restriction enzyme (HaeIII). DNA methylation levels in each sample, which protect HaeIII sites in *AtSN1* from cleavage, were detected by qPCR with primers flanking the sites and normalized using undigested controls. (**D**-**E**) RT-qPCR to quantify the transcript level of (**D**) *AtSN1* under optimum growth conditions and (**E**) *ONSEN* expression in response to 24 h 37°C heat or without treatment (mock). Relative expression in the genotypes indicated below were normalized to the *UBQ10* transcript level as a reference for both *AtSN1* and *ONSEN*. The *ONSEN* assays were further normalized to the Col-0_WT_ mock level. For all qPCR analyses (**C-E**), the error bars represent the standard deviation from the mean (n = 3 technical replicates). (**F**) Model for CLSY-Pol IV docking via the CYC-YPMF motif in the NRPD1 clamp head (asterisk) where there is only one CLSY family member per Pol IV complex. While the locus-specific targeting of RdDM relies on different combinations of CLSY-Pol IV complexes depending on the tissue, as an example, here we show CLSY1 initiating 24 nt siRNA biogenesis and RdDM. At CLSY1-dependent sites Pol IV is recruited using the SHH1 protein, which forms a CLSY1-SHH1 subcomplex that reads H3K9 dimethylation via the Sawadee domain of SHH1.

Next, we compared the effects of the various *nrpd1* mutants and NRPD1-3xF variants on siRNA levels at clusters dependent on different combinations of CLSY proteins. As expected, the two *nrpd1* null mutants and the *clsy quad* mutant have the strongest effects across all categories (**Fig. 4B**). For the NRPD1-3xF_CYC-AAAA_, NRPD1-3xF_AAA-YPMF_, and *nrpd1-50*_YYC-YPMF_ mutants, the siRNA defects were also similar across all categories: NRPD1-3xF_CYC-AAAA_ was always the strongest, NRPD1-3xF_AAA-YPMF_ was always intermediate, and *nrpd1-50*_YYC-YPMF_ was always the weakest (**Fig. 4B**). Thus, these mutations do not appear to selectively affect loci regulated by specific CLSY proteins. However, in the *nrpd1-49*_G72E_ mutant, the *clsy3,4*-dependent clusters were significantly less reduced as compared to the other categories (**Fig. 4B**). In fact, at these clusters, the *nrpd1-49*_G72E_ and *nrpd1-50*_YYC-YPMF_ mutants had similar effects (**Fig. 4B**; red arrow). Assessment of the behavior at *clsy3*- and *clsy4*-dependent siRNA clusters revealed that most of this difference is due to a weaker effect of the *nrpd1-49*_G72E_ mutant on *clsy3*-dependent clusters (**Fig. 4B**; red arrow). These same trends in terms of the overall strengths for each mutant and the selectively weaker effect of the *nrpd1-49*_G72E_ mutant on *clsy3*-dependent clusters are also evident when assessing the normalized read values for each replicate sample across all the 24 nt siRNA categories (**Fig. S7E**). Given the clustering of these mutations in the clamp core and head of NRPD1 (**Fig. 4B**, model inset), which includes the CLSY-docking motif (CYC-YPMF, **Fig. S3A**), these mutant forms of Pol IV are likely all impaired in their association with CLSYs. However, the selective behavior of the *nrpd1-49*_G72E_ mutant on certain Pol IV targets suggests that the location and/or type of *nrpd1* mutation may affect the association of some CLSYs more than others.

To determine the downstream consequences of mutations in the Pol IV clamp head on DNA methylation and TE silencing, the behaviors of the NRPD1-3xF_AAA-YPMF_ and NRPD1-3xF_CYC-AAAA_ variants were compared to WT, *nrpd1-3* null, and *clsy* quad mutant controls. Quantitative Chop-PCR assays, described previously^38^, detected DNA methylation at the *AtSN1* retroelement that is lost in *nrpd1-3* null mutants and rescued after transformation with the NRPD1-3xF_WT_ variant (**Fig. 4C**). Neither the NRPD1-3xF_AAA-YPMF_ nor the NRPD1-3xF_CYC-AAAA_ variant rescued *AtSN1* methylation, as these lines showed methylation levels similar to the *clsy* quad mutant (**Fig. 4C**). In accord with our Chop-PCR data, RT-qPCR experiments detected *AtSN1* transcripts at higher levels in *nrpd1-3* mutants (where DNA methylation is lost) than in WT plants, and *AtSN1* transcripts were reduced in NRPD1-3xF_WT_ lines due to the functional rescue of *nrpd1-3* permitting *AtSN1* silencing. Finally, the higher accumulation of *AtSN1* transcripts in the NRPD1-3xF_AAA-YPMF_ and NRPD1-3xF_CYC-AAAA_ variant lines (**Fig. 4D**) demonstrates that the CLSY-docking motif is required for the silencing of *AtSN1* retroelements.

Previous studies showed that Pol IV plays an important role in silencing LTR/*Copia* retrotransposons of the *ONSEN* family, which are activated by 37°C heat stress^15,38,56,57^. Thus, to assess the dependence of *ONSEN* silencing on the CYC-YPMF motif, we utilized the three NRPD1-3xF variant lines. *ONSEN* transcripts are nearly undetectable by RT-qPCR in Arabidopsis grown under 21/18°C (day/night) conditions (**Fig. 4E**, blue panel). After 24 h of 37°C treatment *ONSEN* accumulates to a much higher level in *nrpd1-3* null plants compared to WT (Col-0_WT_) controls (**Fig. 4E**, orange panel). Moreover, *ONSEN* levels are at WT levels in the NRPD1-3xF_WT_ variant lines but are higher in the NRPD1-3xF_AAA-YPMF_ and NRPD1-3xF_CYC-AAAA_ variant lines, with NRPD1-3xF_AAA-YPMF_ plants showing *ONSEN* levels that phenocopy *clsy* quad mutants (**Fig. 4E**). Together, these findings demonstrate the importance of the CLSY-Pol IV docking motif residues (i.e., CYC-YPMF) for RdDM, for the silencing of *AtSN1* loci, and for the repression of heat-activated *ONSEN* retrotransposons.

## Discussion

Although RNA Pol IV evolved from Pol II, these two polymerases serve very distinct functions: Pol lI transcribes coding regions into mRNAs, while Pol IV works with RDR2 to transcribe TEs and repeats into short dsRNAs that ultimately target DNA methylation and facilitate gene silencing. One critical distinction between these RNA polymerases is how they are recruited to chromatin. Pol II is targeted to defined sequence elements via associations with both specific and general transcription factors, while Pol IV is targeted to distinct chromatin environments via associations with the CLSY family of SNF2-like proteins. Understanding how the structurally-similar Pol II and Pol IV complexes are directed to different targets is critical because the mistargeting of essential genes for silencing by Pol IV or the mistargeting of TEs for expression by Pol II are both dangerous off-target effects. Findings that the CLSYs and RDR2 associate with Pol IV, but with none of the other RNA polymerases^9–12,58,59^, provided the first insights into the unique ability of Pol IV to generate dsRNA. Here we uncovered the mechanism enabling the CLSY proteins to specifically associate with Pol IV and demonstrated this mechanism’s importance in regulating the epigenome.

Specifically, we discovered that NRPD1, a Pol IV-specific subunit, contains a conserved CYC-YPMF motif that is critical for mediating the associations of all four CLSYs with the Pol IV complex and for regulating DNA methylation and siRNA production throughout the genome. In addition, we found that only one CLSY family member associates with a given Pol IV complex. These findings support a “one CLSY per Pol IV” model in which the CLSYs compete for binding to the CYC-YPMF “CLSY-docking” motif to control the distribution of Pol IV across the genome (**Fig. 4F**). Taken together, our work has uncovered a structural innovation within NRPD1 that serves dual roles in controlling DNA methylation patterns. First, it ensures that the CLSYs only interact with Pol IV and second, it creates a highly tunable layer of regulation for Pol IV targeting based the abundances of the CLSYs in each cell. More generally, the separate docking of four CLSYs with Pol IV via a common motif represents a previously unknown mechanism by which SNF2 proteins can, as a group, coordinate noncoding transcription and small RNA biogenesis to maintain genome stability in eukaryotes.

Overall, the identification and investigation of amino acid regions that are exclusively conserved in the largest subunit of Pol IV (as compared to Pol II and Pol V) has provided several major insights into how Pol IV carries out its distinct roles in producing dsRNAs and mediating gene silencing. By applying a probabilistic evolutionary model^60^ to a phylogenetic analysis of ~200 RNA polymerases (**Fig. 1C**, **S1**, **S2A** and **Table S1**), we identified five regions that are exclusively conserved in NRPD1 (EC1-5; **Table S8**). One of these regions, EC2, contains the CYC-YPMF motif that we have now demonstrated acts as a CLSY-docking motif. This fulfils the critical function of ensuring the targeting of Pol IV, but not Pol II, to TEs and repeats throughout the genome. In the recently published cryo-EM structure of the Pol IV-RDR2 complex^26^, two of the other EC regions, EC4 and EC5, were shown to form funnel helices that mediate interactions between RDR2, NRPD1 and its single-stranded RNA product. This Pol IV-RDR2 structural work, the structural analysis of recombinant RDR2^37^ and detailed *in vitro* studies of Pol IV and RDR2 activities^11,25,27,37^ all support a model in which stalling of Pol IV transcription initiates backtracking of the Pol IV complex on the DNA template and threading of its RNA product to the active site of RDR2 via an interpolymerase channel. Thus, as compared to EC2, the EC4 and EC5 regions fulfil a separate, but equally critical, function by ensuring that the synthesis of dsRNA by RDR2 is coupled to transcription by Pol IV, but not by Pol II or Pol V. For the remaining two EC regions, EC1 and EC3, functional data is currently lacking. However, their conservation and positioning within the Pol IV complex suggests testable hypotheses about roles they might play to mediate Pol IV-specific functions. As discussed more below, EC1 is intimately connected with EC2 and thus may also contribute to CLSY binding. For EC3, the position of this loop within the channel connecting Pol IV and RDR2 suggests it could play an important role in guiding the Pol IV transcript into the RDR2 active site.

Determining the locations of conserved regions within the Pol IV-RDR2 structure has also proved informative in assessing their contributions to Pol IV function. By combining an AlphaFold2 prediction of NRPD1 with the Pol IV-RDR2 cryo-EM structure^26^, our resolution of the NRPD1 region corresponding to EC1 and EC2 yielded a series of key insights. First, we found that residues from EC1 and EC2 come together to coordinate a zinc ion within the clamp head region of Pol IV (**Fig. 1F** and **S4D**). Second, we observed that the CYC-YPMF motif of EC2 is exposed at the enzyme exterior on one side of the clamp head (**Fig. 1E**), consistent with its role in mediating Pol IV-specific interactions with the CLSYs. Third, we demonstrated that the other side of the clamp head is lined with positively charged residues predicted to make contact with the downstream DNA template (**Fig. S3D**). Finally, though Pol IV and Pol V fail to copurify with general transcription factors like TFIIA, B, D, E, F or H^9,12,26,61^, comparison of the Pol IV clamp head to the homologous Pol II domain suggests possible activities CLSY proteins could have during Pol IV transcription initiation. The clamp head in Pol II (RPB1 subunit) contacts a Pol II-specific transcription factor TFIIE, which recruits the TFIIH translocase that initially opens downstream DNA^53^. By analogy, a potential mechanism for CLSY function in Pol IV transcription could be to bind the chromatin of transposons, docked to Pol IV via its clamp head, and unwind the downstream DNA. As the timing of the associations between the CLSYs and Pol IV remains unclear, individual CLSYs could bind DNA without Pol IV, followed by Pol IV docking to unwind and transcribe DNA, or they could bind and open DNA prior to Pol IV docking, or the Pol IV complex, already docked with one of the CLSYs, could target and transcribe DNA together as coordinated yet distinct biochemical activities. In **Fig. 4F** we present one plausible version of this process, in which CLSY1-SHH1 recruitment factors facilitate Pol IV transcription into downstream DNA at H3K9me2-marked loci by recruiting Pol IV via its CLSY docking motif.

Notably, the ability of the CYC-YPMF motif to facilitate the association of any one of the four CLSYs with the Pol IV complex distinguishes it from other known Pol IV-specific interactions, which has important consequences for the regulation of DNA methylation patterns. Although RDR2 and SHH1 each have close paralogs in Arabidopsis, only these specific family members associate with the Pol IV complex, demonstrating a high degree of specificity in their binding properties. By comparison, the CYC-YPMF docking site in NRPD1 is more versatile, as it can mediate interactions with all four CLSY partners, suggesting a scenario where the CLSYs compete for binding to Pol IV (**Fig. 4F**). While the rules governing such competition remain to be elucidated, presuming some CLSYs are preferred over others, it would add another dimension to their regulation of DNA methylation. Past studies already demonstrated that the CLSYs are differentially expressed during development and that they target Pol IV to distinct genomic targets^10,20^, resulting in tissue-specific methylation patterns and links between specific CLSYs and several epigenetically regulated traits^16,62^. Within this context, our discovery of a CLSY docking motif in Pol IV suggests a new layer of regulation that could control DNA methylation patterns: competition for binding to the Pol IV complex. Such competition would allow the relative fractions of Pol IV complexes associated with each CLSY to be contextually modulated and also readily reprogrammable. For example, the genomic positions and amounts of DNA methylation could change based not just on the level of each specific CLSY but also on which family members are co-expressed (e.g., Equivalent amounts of CLSY1 expressed with a CLSY that competes better for association with Pol IV versus one that competes worse would lead to different pools of Pol IV complexes, with more Pol IV-CLSY1 complexes in the latter versus the former scenario). Along these same lines, dramatic increases in the expression of a single *CLSY* could rapidly redirect Pol IV by driving the equilibrium of Pol IV composition towards complexes limited to a single CLSY family member. Finally, our discovery that, unlike the mutations in the CYC-YPMF motif that affect siRNA levels similarly regardless of the CLSY responsible, the G72E mutation has a smaller effect on siRNA levels at loci controlled by CLSY3, hints at another possible layer of regulation via natural variation within the NRPD1 subunit of Pol IV. While the aforementioned modes of regulation by the CLSYs remain to be tested, different combinations of these regulatory strategies could explain how a family of just four factors can span the full gamut of DNA methylation regulation—from tuning methylation levels in response to the environment^63^ to generating epigenetic diversity during development^20,22,64^ to germline reprograming during sexual reproduction^23^.

In summary, our findings uncovered a genetic innovation within the Pol IV clamp head (the CYC-YPMF motif) that distinguishes it from other RNA polymerases and enables its targeting to TEs and repeats by mediating interactions with the CLSY proteins. This newly discovered docking location for CLSYs, along with their previously demonstrated targeting functions and the future characterization of their putative chromatin remodeling activities, helps explain how Pol IV functions without the aid of TFII general transcription factors that are critical for Pol II recruitment and initiation. Given the existence of other specialized factors that enable RNA polymerases to transcribe in heterochromatin, like the TFIIA-related Moonshiner protein in Drosophila that directs Pol II to initiate piRNA production^13^, our evolutionary rate and structural modeling approaches could provide further insights into atypical Pol II recruitment factors, or more broadly, into protein families with specialized activities in other processes and species.

## Materials and Methods

### Plant materials

#### Genetic mutants

Previously published Arabidopsis T-DNA insertion mutants used in this study include: *nrpd1-3* (SALK_128428)^65^, *nrpd1-4* (SALK_083051)^66^, *clsy1-7* (SALK_018319)^67^, *clsy2-1* (GABI-Kat line 554E02), *clsy3-1* (SALK_040366) and *clsy4-1* (SALK_003876)^10^. The *nrpd1-49*, *nrpd1-50* and *nrpd1-51* point mutations were described by Ferrafiat et al. 2019^38^.

#### NRPD1 epitope tagged lines

The *nrpd1-3* null mutant was transformed using Agrobacterium with different *NRPD1p::NRPD1-3xF* (BastaR) constructs that express either WT (NRPD1-3xF_WT_) or mutant (NRPD1-3xF_AAA-YPMF_ or NRPD1-3xF_CYC-AAA_) forms of the NRPD1-3xF protein. Basta resistant T_2_ progeny homozygous for the *nrpd1-3* mutation in which the NRPD1-3xF protein was strongly expressed (**Fig. S5A**) were selected for IP-MS experiments. In separate experiments, these lines were super-transformed with a *CLSY1p::3xHA-CLSY1* (HygR) construct and the resulting T_2_ progeny were selected for co-IP and IP-MS experiments based on hygromycin resistance as well as the accumulation of the 3xHA-CLSY1 and NRPD1-3xF proteins.

#### RDR2, SHH1 and CLSY epitope tagged lines

Wild-type Col-0 plants were transformed using Agrobacterium with *35S::RDR2-eGFP* or *35S::SHH1-3xHA* constructs. T_2_ progeny were selected based on hygromycin resistance and detection of the RDR2-eGFP or SHH1-3xHA proteins. T_2_ lines were crossed with the NRPD1-3xF_WT_, NRPD1-3xF_AAA-YPMF_, or NRPD1-3xF_CYC-AAA_ lines and F_1_ progeny were used for co-IP experiments.

All the N- and C-terminal 3xF tagged CLSY1, CLSY3 and CLSY4 lines used here are in their respective *clsy* mutant backgrounds and are driven by their respective endogenous promoters. Four lines were previously characterized and shown to complement their respective *clsy* mutant phenotypes by Zhou et al. 2022^20^ and the remaining lines were characterized in this study (**Fig. S6**). In all cases, T_3_ lines homozygous for the various 3xF-tagged CLSYs were identified based on drug selection using Hygromycin and seeds from subsequent generations were grown directly on soil under Salk greenhouse conditions for the IP-MS experiments. For the Co-IPs between CLSY3 and either the NRPD1-3xF_WT_ or NRPD1-3xF_CYC-AAAA_ lines, a different CLSY3 tagged line was used that contains both a 3xF epitope and a Biotin Ligase Recognition Peptide (CLSY3-3xF-BLRP) that once biotinylated can be captured using streptavidin beads. This construct, which is also driven by the endogenous *CLSY3* promoter and complements the *clsy3* mutation^20^, was crossed to the NRPD1-3xF_WT_ or NRPD1-3xF_CYC-AAAA_ lines and F_1_ progeny that were double drug selected (Hygromycin and Basta) were used for the co-IP experiments (**Fig. 2E** and **S5D**).

### Antibodies

The largest subunit of Arabidopsis Pol IV was detected using a native antibody against a peptide from the AtNRPD1 C-terminus. The second largest subunit of Pol IV was detected using an antibody against a peptide from the AtNRPD2 N-terminus (see Ferrafiat et al. 2019). The various 3xF tagged NRPD1 and CLSY proteins were detected using a monoclonal anti-FLAG-HRP antibody (Sigma #A8592). RDR2-eGFP was detected using an anti-eGFP polyclonal antibody^68^, whereas 3xHA-CLSY1 and SHH1-3xHA were detected using a monoclonal anti-HA-HRP antibody (Sigma #H6533).

### Molecular cloning and complementation assays

Cloning of the N- and C-terminal 3xF and 3xF-BLRP CLSY lines are as previously described in Zhou et al. 2022^20^. For all other *A. thaliana* transgenic lines, the plant transformation vectors used were generated via the Multisite Gateway approach (ThermoFisher Scientific). Genomic sequences and promoters were amplified by PCR from Col-0 genomic DNA with primers that flanked the 5’-end Multisite Gateway sites (**Table S6**). The amplified sequence was cloned into the pDONR221 vector and the promoter was cloned into pDONRP4p1r using BP Clonase II (ThermoFisher Scientific). To obtain NRPD1 motif mutant lines, Gibson Assembly (NEB) was performed using a WT *A. thaliana* genomic NRPD1 fragment that we cloned into pDONR221 via BP Clonase II reaction (pENTR-NRPD1_WT). Then, an LR Clonase II reaction was performed to insert the *NRPD1p::NRPD1-3xF* sequence into the pB7m34GW plant expression vector (BastaR), or to insert the SHH1p::SHH1-3xHA, RDRp::RDR2-eGFP or CLSY1p::3xHA-CLSY1 sequences into pH7m34GW vector (HygR). The expression vectors were validated by sequencing before plant transformation using *Agrobacterium tumefaciens* GV3101.

For the 3xF tagged CLSY lines used in **Fig. 3A** and **B**, complementation was as assessed by genome-wide smRNA-seq experiments and visualized as volcano plots as described in Zhou et al. 2022^20^. In addition to the lines published in Zhou et al. 2022^20^, which correspond to CLSY1 Line1 (ins#1 in Zhou et al. 2022^20^), CLSY3 Line 2 (ins#3 in Zhou et al. 2022^20^), and CLSY4 Line 2 (ins#1 in Zhou et al. 2022^20^), CLSY1 Line2, CLSY3 Line 1, and CLSY4 Line 1 other lines were sequenced and process in parallel and thus can be compared to the previously published controls as shown in **Fig. S5A-C**. The coverage, mapping, and size distributions of these new smRNA-seq samples are included in **Table S5**.

### Phylogenetic and exclusive conservation analyses

The amino acid sequences from the largest subunits of the three DNA-dependent RNA polymerases (Pol II, Pol IV, and Pol V) were obtained from Phytozome (https://phytozome-next.jgi.doe.gov/) and the National Center for Biotechnology Information (NCBI) using BLASTP with the queries AtNRPB1, AtNRPD1 or AtNRPE1, respectively. In total, 202 sequences from 56 species were analyzed in Geneious (v11.1.5, https://www.geneious.com/). All multiple sequence alignments were performed using MUSCLE (v3.8.425, default parameters). Phylogenetic Trees were built with Geneious Tree Builder using default parameters. Conservation scores were calculated using the ConSurf analysis tool, with AtNRPD1 as the reference sequence and the default settings ^49^. AtNRPD1 conservation among the plant NRPD1s was based on a MUSCLE alignment of 58 protein sequences with NRPD1 domain features. The positional conservation in AtNRPD1 was calculated via comparison to all NRPB1, NRPD1, or NRPE1 subunits using an alignment of the plant sequences recognizable as Pol II, Pol IV, or Pol V largest subunits (**Table S1**) but removing the 12 bryophyte proteins in NRPD1/E1-like subclades (final dataset: 190 proteins from 52 species). To quantify exclusive conservation (EC) at individual AtNRPD1 sites, we took the NRPD1 conservation score and subtracted the maximum of NRPB1 and NRPE1 conservation scores at each position. A high EC_NRPD1_ score represents an amino acid specific to NRPD1, whereas a low EC_NRPD1_ indicates that the amino acid there is similar to at least NRPB1 or NRPE1 (or specific to one of these). In order to identify exclusively conserved regions (EC1, EC2, EC3 …), a centered moving average (k=9) was applied to EC_NRPD1_. Positions with averages ≥ 2 were found, then all positive consecutive values surrounding these seed positions were selected and expanded to include positions with positive values separated by < 9 aa. These EC_NRPD1_ scores and hotspots were then plotted using R (**Fig. 1D**). C-terminal domains (CTDs) of the NRPB1, NRPD1 and NRPE1 subunits were masked for the ConSurf plot, because NRPB1 and NRPE1 have highly repetitive and divergent CTDs that do not show the consistent alignment of homologous positions for all species.

### Structural analysis

All structural analyses were performed in ChimeraX v1.3 to v1.6^69^. Inspection of the cryo-EM density from AtNRPD1 in Pol IV-RDR2 (EMD-31305) after Gaussian filtering revealed the presence of a globular domain corresponding to an unmodelled portion (aa 93-203) of the atomic model 7EU0. The structure of the unmodelled Pol IV domain was predicted using the AlphaFold2 online service (https://www.alphafold.ebi.ac.uk/) and UniProt ID (Q9LQ02) for AtNRPD1^70^. The AlphaFold2 model was aligned with the AtNRPD1 model 7EU0 using Matchmaker^71^. The CYC-YPMF domain of interest in the predicted model was then rigid body fitted to the experimental cryo-EM density. The pLDDT score combined with the proper fit of the domain in the density and the presence of a similarly positioned domain in Pol II structures of human (6GMH) and yeast (7O75) confirmed the proper attribution of the density to this unmodelled portion of AtNRPD1 (**Fig. S3A, B, C**). The AlphaFold2-predicted structure (**Fig. 1E**, pink residues) overlaps amino acids that have been previously assigned to a putative Pol IV ‘clamp head’ (aa 85-219)^26^, again based on comparison to Pol II structures.

The AtNRPD1 residues Cys97, Cys100, Cys118 and Cys121 were identified as a putative zinc binding site using the online Zincbindpredict tool^54^ and supported by the AlphaFill software also predicting a zinc ion at this site^55^. As a positive control, another zinc binding site the AtNRPD1, composed of residues Cys56, Cys59, Cys67, and His70, was identified by Zincbindpredict. These residues are conserved in humans and yeast and are known to coordinate zinc^50^. For **Fig. S4A**, the atomic model was colored according to the ConSurf analysis using the command “color byattribute” and an attribute file containing the centered moving average of NRPD1 exclusive conservation (EC_NRPD1_) values. The NRPD1 model in **Fig. S4A** displays this moving average as a continuous color gradient ranging from ≥0 (white) to 3.5 (purple). Discrete colors were assigned in **Fig. S4B** to the five NRPD1 regions with EC_NRPD1_ peaks (EC1-EC5) shown in **Fig. 1D**. The electrostatic potential analysis was performed using the “coulombic” command^69^.

### Denaturing protein extraction and western blot analyses

Samples for steady-state protein analyses (**Fig. S5A**) were obtained using a phenol-based extraction on frozen powder from *A. thaliana* flowers^72^. The resulting pellets were resuspended in resuspension buffer (RB; **Table S7**) and proteins levels were quantified by Lowry (Bio-Rad #5000113, #5000114, #5000114). 600 µg aliquots of each sample were pre-heated at 95°C for 5 min. and then loaded on a 6% polyacrylamide gel. After SDS-PAGE migration and western blot transfer to Immobilon-P membrane (Millipore IPVH00010), the membrane was incubated with 5X blocking solution (0.75 g powder milk in 15 mL 1X PBS, 0.1% Tween20) for 30 min. and then overnight with the primary antibody at 4°C using antibody-specific dilutions (anti-FLAG-HRP, 1:15000; anti-HA-HRP, 1:15000; anti-eGFP, 1:20000; or Anti-NRPD2, 1:2500). Next, the membrane was washed with 15 mL of 1X PBS, 0.1% Tween20 for 20 min., three times each at room temperature and incubated with a secondary antibody coupled to horseradish peroxidase (1:15000) for 2 h at 4°C, followed by chemiluminescent detection on a Fusion FX7 Edge (Vilber) using Lumi-Light Plus Substrate (Roche). The membrane was then stripped with Restore PLUS Western Blot Stripping Buffer (Thermo Scientific), washed, blocked and incubated with the next antibody.

### Affinity purification of Pol IV complexes

#### NRPD1-3xF, 3xHA-CLSY1, SHH1-3xHA and RDR2-GFP

Flower tissue of *A. thaliana* was ground in liquid nitrogen. 1.5 g of flower powder was then mixed with 3 mL of lysis buffer (LB; **Table S7**) at 4°C for 20 min. The resulting extracts were centrifugated twice at 16,000 rcf at 4°C for 15 min. to obtain clear supernatants. For NRPD1-3xF and 3xHA-CLSY1 IP-MS, the same protocol was used. Each supernatant was incubated with 50 µL of either anti-FLAG or anti-HA antibodies associated with magnetic beads from the corresponding Miltenyi Biotec Isolation kit on a wheel at 10 rpm at 4°C for 45 min. µMACS DYKDDDK Isolation Kits were used for anti-FLAG IPs and HA Isolation Kits for anti-HA immunopurification (Miltenyi Biotec #130-101-591 and #130-091-122). Next, each µColumn (Miltenyi Biotec #130-042-701) was conditioned with 400 µL of lysis buffer. Magnetic bead-incubated samples were loaded in 400 µL batches on the Miltenyi stand. These columns were washed with 4 x 400 µL of wash buffer (WB; **Table S7**), followed by 200 µL of µMACS kit wash buffer (Miltenyi Biotec #130-042-701). The protein complexes were then eluted into 1.5 mL Eppendorf tubes using 3 x 35 µL of pre-heated 95°C elution buffer (Miltenyi Biotec #130-042-701). The samples were heated to 95°C for 5 min. and stored at −20°C prior to MS analysis at the Strasbourg-Esplanade Proteomics Facility. For the co-IP experiments shown in **Fig. 2F, G** and **H**, the same IP protocol was followed, except that in addition to the abovementioned kits, the Miltenyi anti-GFP Isolation Kit was used for anti-GFP IPs (Miltenyi Biotec #130-091-125), and eluted protein samples were analyzed by western blot analysis, as described in Ferrafiat et al. 2019^38^.

#### N- and C-terminal CLSY 3xF IPs

The affinity purification was performed as described in Law et al. 2011^9^ using 10 g of Arabidopsis flowers with the following modifications. The IP buffer was supplemented with 10 mM bortezomib (IB, **Table S7**), anti-Flag M2 magnetic beads (Sigma #M8823) were used, and after five bead washes with 1000 μl of fresh IP buffer, the proteins were eluted twice (15 min. at RT) by incubation with 3x-FLAG peptide (Sigma #F4799) at a concentration of 100 mg/mL in PBS buffer. The eluted samples were precipitated using the Trichloro Acetic Acid (TCA) method and heat-dried pellets were stored at 4°C until Mass Spectrometry analysis.

### Liquid chromatography-tandem mass spectrometry proteomics for 3xF-tagged NRPD1 variants and 3xHA-tagged CLSY1

After Pol IV complex immunoprecipitation, the frozen protein samples were transferred to the Strasbourg-Esplanade Proteomics Facility at the Institut de Biologie Moléculaire et Cellulaire (IBMC). Proteins were precipitated by adding 5 volumes of glacial 0.1 M ammonium acetate in 100% methanol, then stored for 12 h at −20°C, washed with 0.1 M ammonium acetate in 80% methanol and dried under vacuum. The dry pellets were resuspended in 50 mM ammonium bicarbonate, reduced with 5 mM dithiothreitol (DTT) for 10 min. at 95°C and alkylated with 10 mM iodoacetamide for 30 min. at room temperature in the dark. The resulting proteins with carbamidomethyl groups on their cysteines were digested with 300 ng of sequencing-grade porcine trypsin (Promega, Fitchburg, MA, USA) and injected on an Easy-nanoLC-1000 system coupled to a Q-Exactive+ mass spectrometer (Thermo Fisher Scientific, Germany) using a data-dependent acquisition strategy with 160 min. gradients. Arabidopsis proteins were identified via comparison to the TAIR10 database (27,222 protein sequences, https://www.arabidopsis.org/), analyzed using the Mascot algorithm (version 2.6.2, Matrix Science) and spectral counts were validated in Proline software (v2.0, ProFI) with analysis parameters that accommodate a false discovery rate of <1% at the peptide spectrum matches and protein levels. Once Arabidopsis protein IDs were determined, statistic comparisons of the spectral counts in different samples were carried out to obtain, after a DESeq2 normalization, p-values based on the negative binomial distribution (R script IPinquiry_v04) (https://github.com/hzuber67/IPinquiry4) and adjusted by the Benjamini-Hochberg method. The raw data for these IP-MS analyses are available via ProteomeXchange with the identifier PXD047743.

### Proteomic characterization of affinity purified 3xF-tagged CLSY complexes

For all four batches of IP-MS experiments (see **Fig. 3B** and **Table S3**), acetone precipitated protein pellets obtained after immunoprecipitation of the CLSY proteins were resuspended in 4M urea, 100mM Tris-Cl, pH 8.5. This was followed by reduction and alkylation by the sequential addition of 5 mM tris(2-carboxyethyl)phosphine and 10 mM iodoacetamide. Reduced and alkylated samples were diluted to reduce urea concentration to 2M followed by proteolytic digestion with Lys-C and trypsin at 37°C overnight. The digested peptides were subjected to offline SP3-based peptide clean-up^73^ and subsequently analyzed by LC-MS/MS. Briefly, peptides were separated by reversed phase chromatography using 75 μm inner diameter fritted fused silica capillary column packed in-house to a length of 25 cm with bulk 1.9 mM ReproSil-Pur beads with 120 Å pores^74^. For batches 1 and 2, the samples were analyzed on a Thermo Fisher Fusion Lumos mass spectrometer coupled to a Dionex Ultimate 3000 UHPLC using a data-dependent acquisition strategy with an MS1 resolution (r) of 120K followed by sequential MS2 scans at a resolution of 15K^74^. For batch 3, the samples were analyzed using a Thermo Fisher Q-Exactive coupled to an easyLC 1000 using a data-dependent acquisition strategy as previously described^75^. For batch 4, the samples were analyzed on Thermo Fisher Fusion Lumos mass spectrometer coupled to a Dionex Ultimate 3000 UHPLC using a data-independent acquisition strategy in which 90 variable isolation windows (resolution = 15K) were employed to collect MS/MS spectra across a 400 – 1600 m/z range (vDIA)^76^. For batches 1-3, the data generated by LC-MS/MS were analyzed using the MaxQuant bioinformatic pipeline^77^. The Andromeda integrated in MaxQuant was employed as the peptide search engine and the data were searched against the Arabidopsis database (Uniprot Reference UP000006548). Briefly, a maximum of two missed cleavages was allowed. The maximum false discovery rate for peptide and protein was specified as 0.01. Label-free quantification (LFQ) was enabled with LFQ minimum ratio count of 1. The parent and peptide ion search tolerances were set as 20 and 4.5 ppm respectively. For batch 4, the vDIA data was analyzed using DIA-Umpire to generate pseudo-MS2 spectra from DIA data followed by MSFragger for database searching^78,79^. The MaxQuant and MSFragger output files were subsequently processed for statistical analysis of differentially enriched proteins using the R package IPinquiry4 (https://github.com/hzuber67/IPinquiry4). The raw data for all four batches of IP-MS data are available through the MassIVE repository via the identifier MSV000093500.

### Mass spectrometry data visualization

The data visualization was done in R (v4.2.3) using the ggplot2^80^ and ggpubr^81^ libraries. The results from the IPinquiry Analysis were loaded and volcano plots were made using the produced Log_2_FC and adjusted p-values, highlighting the cofactors and core Pol IV proteins. The same dataset was used to produce a variety of balloon plots using the spectral count and adjusted p-values respectively as size and color factors.

### CLSY3-NRPD1 co-IPs

For each genotype, flower tissue (stage 12 and younger) was collected, flash-frozen and ground to a fine powder in liquid nitrogen. For each IP, 0.15 or 0.20 g of powder per genotype was resuspended in two volumes of IP buffer (IB; **Table S7**), spun twice (max speed for 10 min. at 4°C), and the supernatant was added to a new tube with 75 μL of IP buffer-washed magnetic Streptavidin beads (M-280 Dynabeads #11205D). The sample and bead solutions were rotated (30 min. at 4°C) and unbounded proteins were removed by five washes in 1000 μl of fresh IP buffer (2 min. at 4°C). The proteins were eluted by heating the beads (95°C for 5 min.) in 70 μL of BME 4X Laemmli buffer (Biorad #1610747). Prior to loading on a 7.5% TGX SDS-PAGE gel (Biorad #5671024), 30 µL of each sample was heated (5 min. at 95°C) and spun down (max speed for 3 min. at RT). The proteins were separated at RT in running buffer (RB; **Table S7**) for 30 min. at 60 V followed by 1 hr. 30 min. at 150 V. Proteins were transferred onto a 0.45 μm PVDF Amersham Hybond membrane (#10600023) at 4°C in transfer buffer (TB; **Table S7**) for 1 hr. 30 min. at 160 mA. The membrane was blocked in TBS-T containing 3% BSA for 1 hr. 30 min. at RT and incubated overnight with a monoclonal Anti-FLAG antibody coupled to horseradish peroxidase (Sigma #A8592; dilution 1:10,000 in TBS-T containing 3% BSA). After five, 5 min. rinses in wash buffer (WWB; **Table S7**) at RT under slow agitation (18 rpm), the membrane was incubated with Pierce ECL2 Western Blotting substrate (#PI80196X3) for 5 min. in the dark. The Azur Sapphire system was used for the chemiluminescence imaging.

### RNA extraction and RT-qPCR

Total RNA from Arabidopsis flower tissue was extracted with TRIzol reagent, treated with DNase I (ThermoFisher Scientific #EN0521) and purified using phenol-chloroform and ethanol precipitation. The reverse transcription was performed using SuperScript IV Reverse Transcriptase (Invitrogen #18090050) and RiboLock RNase Inhibitor (ThermoFisher Scientific #EO0381) with 1 μg of RNA previously treated with DNase. The cDNA obtained is analyzed by qPCR LightCycler 480 II (Roche) using Takyon No ROX SYBR 2X (Eurogentec #UF-NSMT-B0701) and primers to detect *ONSEN* and *AtSN1* transcripts.

### Small RNA blot

Small RNA isolation from Arabidopsis flowers was performed as described in Böhrer et al. 2020^82^. In short, total RNA was extracted using TRIzol and then size-fractionated using the RNA clean-up protocol of a RNeasy Midi kit (Qiagen #75144). Once the samples are dehydrated using SpeedVac (SPD111V, ThermoFisher Scientific), they are loaded onto 16% polyacrylamide gel and run for 1 h at 15W. The RNAs are transferred into Hybond-N+ nylon membrane (GE Healthcare #RPN203B) at 300 mA for 2 h at 4°C and fixed by UV at 1400J. The membrane is washed with 50 mL 2X SSC (Saline sodium citrate), blocked for 3h with 20 mL of PerfectHyb Plus Hybridization (Sigma-Aldrich #H7033-1L), and incubated overnight between 35-40°C, depending on the probe. The PNK and Klenow probes preparation are described in Böhrer et al. 2020^82^. The membrane is washed three times with 20 mL of 0.5% SDS, 2X SSC and the membrane is exposed to a phospho-imager screen into a cassette for seven days. The signal of the screen is revealed by phospho-imager (Amersham Typhoon, GE Healthcare). To rehybridize the membrane, the probe was stripped with boiled 0.1% SDS twice for 20 min., and incubated with 20 mL of PerfectHyb Plus Hybridization with the new probe.

### Small RNA data processing

All samples included in **Table S5** were processed as follows. Raw reads were trimmed using cutadapt (v1.18)^83^ to remove adapters (-a AACTGTAGGCACCATCAAT) and reads shorter than 15 nt (-m 15). Trimmed reads were mapped to the Arabidopsis TAIR10 genome using ShortStack (v3.8.5)^84^ allowing one mismatch (--mismatches 1) and allowing multi-mapped reads that will be guided to a specific location with the fractional-seeded algorithm (-mmap f). To meet the characteristics of Pol IV transcription^28^, the mapped reads were further filtered to keep only perfectly-mapped reads or reads with a single mismatch at the 3’-ends using the filter function of bamtools (v2.5.1)^85^ together with a previously published script JSON_findPerfectMatches_and_TerminalMisMatches_v3^10^. To facilitate small RNA quantification, Tag Directories were generated from the filtered bam files using the makeTagDirectory function of HOMER (v4.10)^86^ with the following options: -format sam -mis 1 -keepAll. Split Tag Directories with a certain smRNA size (21-24 nt) were made using a previously published perl script splitTagDirectoryByLength.dev2.pl^10^.

### Core small RNA cluster comparison

To validate the use of the previously reported master set of small RNA clusters (n=12,939; Zhou et al. 2022^20^), we calculated how many 24 nt siRNAs from the current dataset and the Ferrafiat et al. 2019 dataset were covered by these clusters. To count the number of 24 nt siRNAs covered by the master set of small RNA cluster, we first used BBmap to filter the bam files to retain reads that are 24 nt in length (reformat.sh minlength=24 maxlength=24) and then keep reads that are located within specific regions using samtools^87^. The pie charts showing the 24 nt siRNAs covered by each of the categories were plotted in R (https://www.r-project.org/).

### Differential expression analysis

To identify differentially expressed 24 nt siRNA clusters compared to the WT controls, DESeq2^88^ was used to perform the differential analysis. As 24 nt siRNAs in some of the RdDM mutants (i.e., *pol iv* mutants) are dramatically decreased, siRNAs of this size class cannot be used to estimate the library size. Instead, we used all reads that are mapped to the TAIR10 genome and also meet the criteria of the filtering process (see the Small RNA data processing section) to calculate the library size factors. Differentially expressed siRNA clusters were identified with fold change (FC) ≥ 2, which is equivalent to |log_2_FC| ≥ 1, and a false discovery rate (FDR) < 0.01.

### Visualization of 24 nt siRNAs

Normalized 24 nt siRNAs at the master set of small RNA cluster was generated using the annotatePeaks.pl function of HOMER^86^ with the “-size given -fpkm -len 1” options. Violin plots comparing 24 nt siRNA levels across different sets of clusters were made using the ggplot2^80^ package of R (https://www.r-project.org/). Half violin plots and volcano plots were made using the ggplot2^80^ package of R (https://www.r-project.org/) based on the DESeq results.

### DNA methylation detection

Genomic DNA extraction was conducted using the Nucleon Phytopure Kit (Cytiva #RPN8511), treated with RNase A/T1 (2 mg/mL) and purified using phenol-chloroform and ethanol precipitation. 500ng of DNA were digested with methylation-sensitive HaeIII restriction enzyme and qPCR was conducted to amplify undigested DNA with primers flanking the HaeIII sites in the *AtSN1* retroelement (**Table S6**).

### Heat stress and retrotransposon detection

Seeds were sterilized with 70% ethanol and 4% bleach and grown on solid 0.5X MS medium (Murashige & Skoog, M0255, Duchefa) (1% sucrose, agar, pH 5.7) under long-day conditions (16 h light) at 21°C and short night conditions (8 h) at 17°C. 14-day-old plants were incubated in liquid MS medium under control stress (24 h at 21°C) or heat stress (24 h at 37°C). An RNA extraction was conducted following the protocol in Böhrer et al. 2020^82^ and qPCR was executed with transcript-specific primers (**Table S6**) to measure *ONSEN* copy numbers was executed as described in Ito et al. 2011^15^ and Thieme et al. 2017^57^.

## Acknowledgements

We thank lab members and colleagues for helpful comments and fruitful discussions. The Q-Exactive+ mass-spec instrumentation was funded by the IdEx Equipement mi-lourd grant (University of Strasbourg 2015). J.W. was supported by the NIH (GM089778). F.W. was supported by the Swiss National Science Foundation with a Swiss Postdoc Fellowship (project 210561). J.L. was supported by the NIH (GM112966). G.X. was supported by a postdoctoral fellowship from the Paul F. Glenn Center for Biology of Aging Research at the Salk Institute. L.M. was supported by the Catharina Foundation postdoctoral fellowship. This work was also supported by the NGS Core Facility of the Salk Institute with funding from NIH-NCI CCSG: P30 CA01495, NIH-NlA San Diego Nathan Shock Center P30 AG068635, the Chapman Foundation and the Helmsley Charitable Trust. T.B. was supported by the ANR (ANR-17-CE20-0004), and by the Interdisciplinary Thematic Institute IMCBio (ITI 2021-2028 program), including funds from IdEx University of Strasbourg (ANR-10-IDEX-0002), SFRI-STRAT’US (ANR 20-SFRI-0012) and EUR IMCBio (ANR-17-EURE-0023) in the framework of the French Investments for the Future Program.

## Author contributions

Protein extractions and IPs were performed by L.F., B.R., L.M., P.H., C.H., M.Z. and J.E. Molecular cloning and transgenic lines were generated by B.R., L.F., M.Z., L.M., J.E., C.C. Proteomics analyses were conducted by V.P., J.C., L.K., J.W., P.H. Structural modelling was done by F.W. Bioinformatics and phylogenetics analyses were performed by G.X., C.M., B.R., L.F., C.C. and T.B. The manuscript was written by B.R., L.F., L.M., J.L. and T.B.

## SUPPLEMENTARY DATA

### Supplementary figure legends

**Figure S1.**
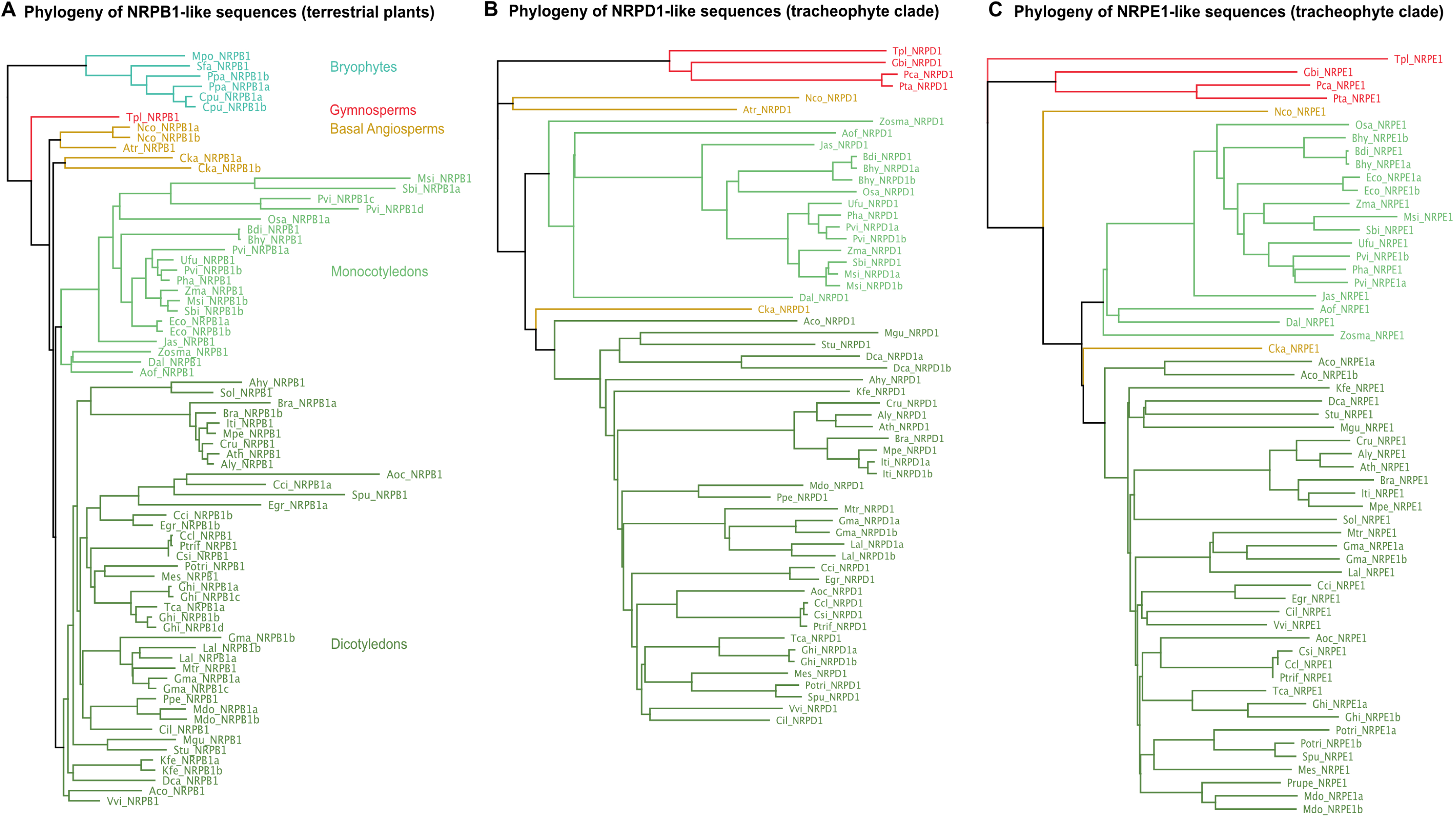
Phylogenetic analyses of the largest subunits of Pol II, Pol IV and Pol V in terrestrial plants. Phylogenetic tree representing available **(A)** NRPB1, **(B)** NRPD1 and **(C)** NRPE1 proteins from diverse terrestrial plants. The tree covers terrestrial plant species including gymnosperms and diverse angiosperms (basal, monocotyledons, and dicotyledons). The protein sequences were obtained from the Phytozome13 and NCBI databases. The phylogenetic analyses were performed with the Geneious software package using MUSCLE for multiple sequence alignment and the Geneious neighbor-joining tree builder with the Jukes-Cantor distance model. The full list of RNA polymerase subunits used in this analysis (including the corresponding species names, database accessions and protein characteristics) is provided in **Table S1**.

**Figure S2.**
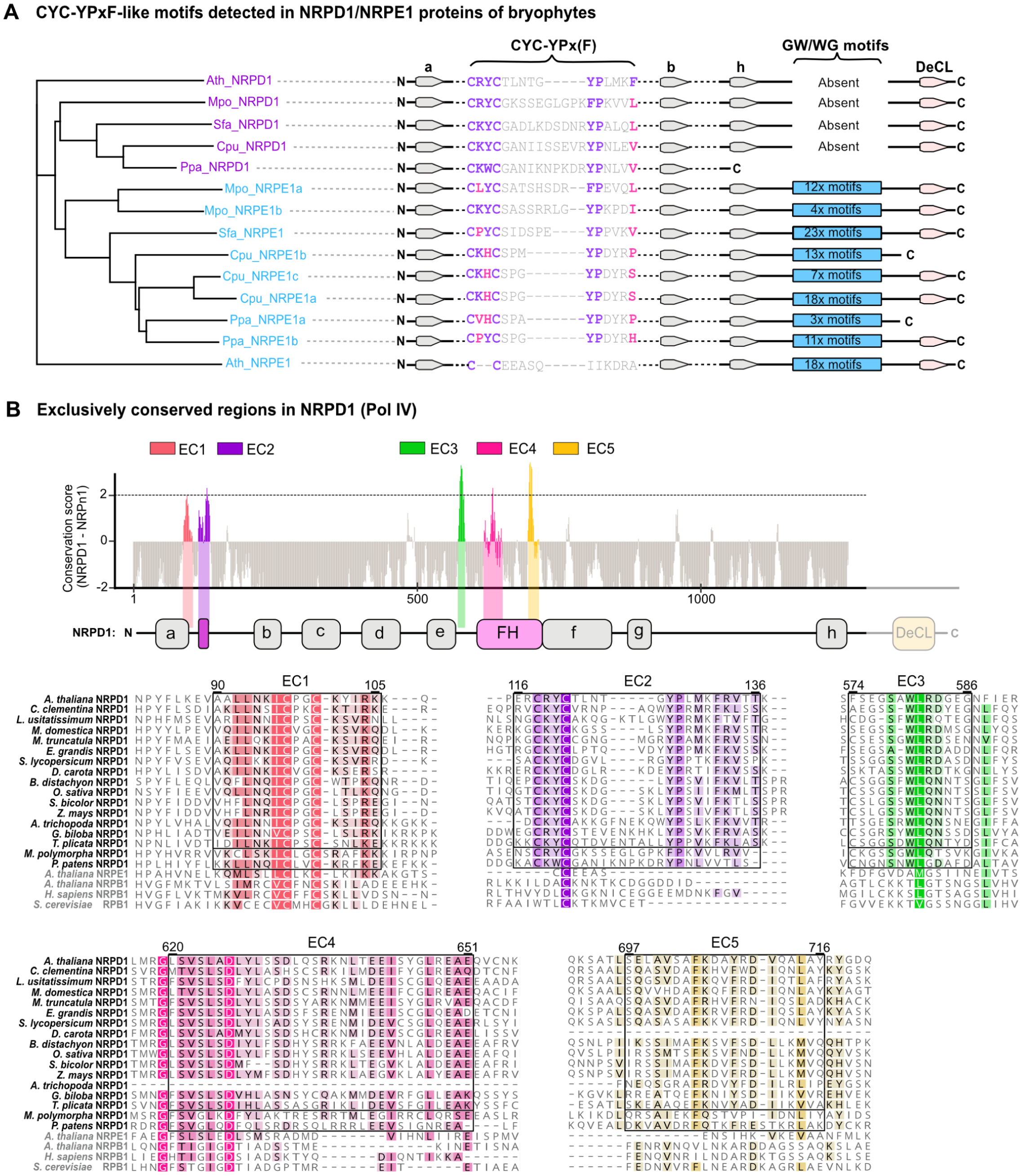
Bryophyte NRPD1 and NRPE1-like proteins have divergent CYC-YPxF motifs as compared to NRPD1 in vascular plants. **(A)** Phylogenetic tree of the CYC-YPxF region in bryophyte NRPD1 and NRPE1-like proteins compared to corresponding *A. thaliana* orthologs. Putative bryophyte NRPD1 subunits (Pol IV) are indicated in purple and putative NRPE1 subunits (Pol V) are indicated in light blue. Bryophyte NRPE1-like protein identities were inferred in this analysis based on GW/WG repeat sequences found in the C-terminal domain (CTD) of each protein. (**B**) ***top panel:*** Reproduction of the Fig. 1D. ConSurf analysis of NRPD1 proteins from 52 plant species compared to NRPB1 and NRPE1. The five protein regions that are peaks of NRPD1-exclusive conservation (labeled EC1-EC5) are plotted in color along the subunit’s domain architecture. ***bottom panels:*** Exclusively conserved (EC) regions are marked by boxes in the multiple sequence alignments of NRPD1 proteins of diverse species, with conserved residues in colors matching the distinct EC regions. The amino acid numbering above the boxed EC regions is based on NRPD1 of *A. thaliana*. Corresponding NRPD1 regions from two representative bryophytes, *Marchantia polymorpha* and *Physcomitrium patens*, are boxed separately to show how the Pol IV-specific motifs have diverged in non-vascular plants. Positions and domains with low NRPD1 exclusive conservation are in grey.

**Figure S3.**
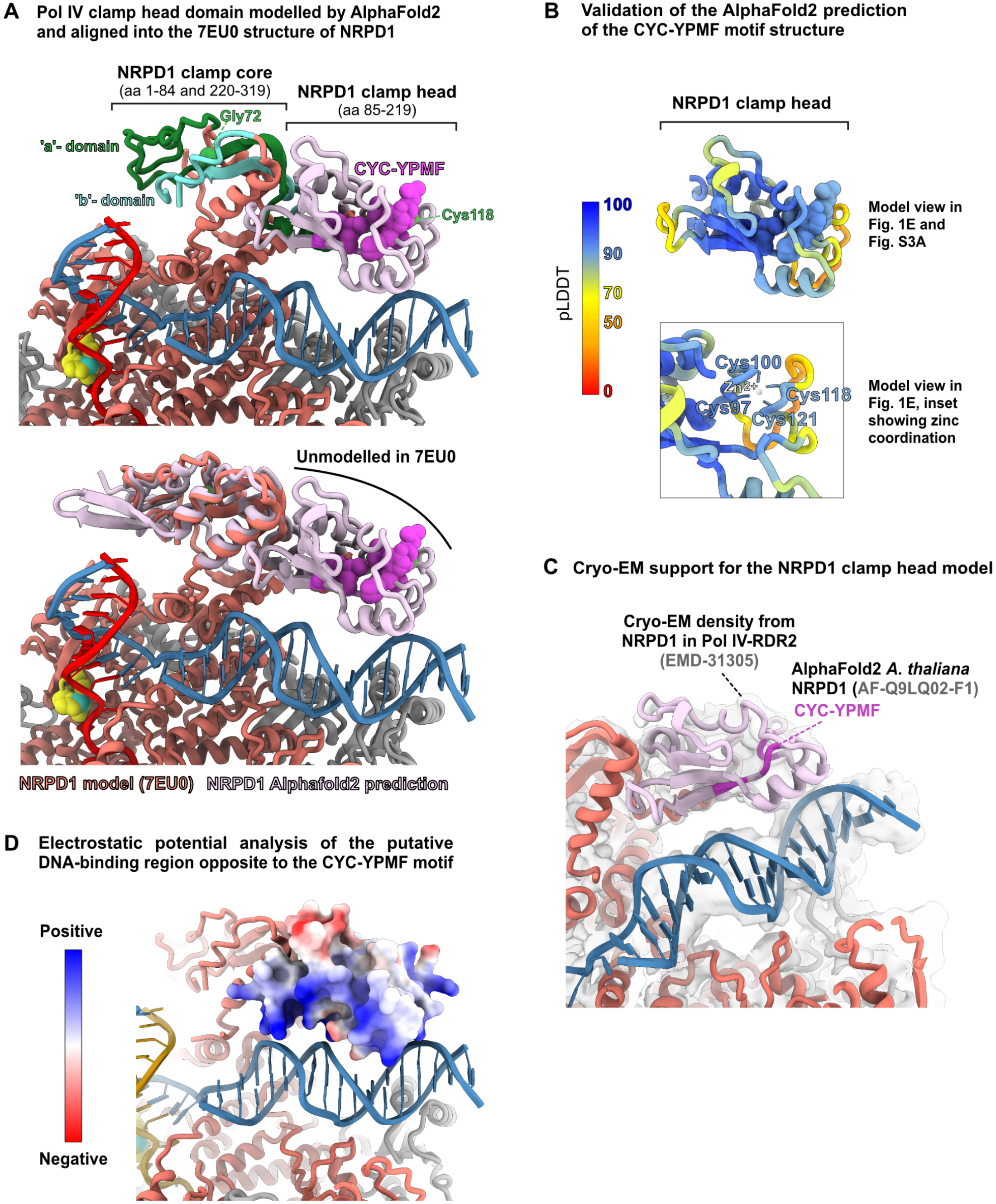
Structural analysis and validation of the Pol IV clamp head model. **(A)** AlphaFold2 prediction of the previously unmodelled ‘clamp head’ domain structure of NRPD1 that contains the CYC-YPMF residues (bright purple) and is located close to the DNA template (blue). ***top panel:*** an AlphaFold2 NRPD1 ‘clamp head’ prediction was fit into the NRPD1 ‘clamp core’ structure to form an extended Pol IV model: aa 1-84 and 220-319 (clamp core) derive from the 7EU0 structure of NRPD1, while aa 85-219 are from the AlphaFold2 prediction of NRPD1 (AF-Q9LQ02-F1) and correspond to the clamp head (light pink). The NRPD1 positions Gly72 and Cys118 (green) are mutated in the *nrpd1-49* and *nrpd1-50* mutants, respectively (Ferrafiat et al. 2019). ***bottom panel:*** the 7EU0 structure of NRPD1 (coral) is superimposed with the AlphaFold2 prediction of the full NRPD1 clamp (pink). **(B) *top panel:*** The CYC-YPMF-containing clamp head domain is shown as predicted by AlphaFold2 with colors representing AlphaFold2’s per-residue confidence score (pLDDT). ***bottom panel:*** pLDDT for the AlphaFold2 clamp head prediction including the Cys97-Cys100-Cys118-Cys121 residues coordinating a zinc and with aa 166-172 removed to avoid obscuring the view of the cysteine side chains. Residues with pLDDT >90 are expected to be modelled to high accuracy, scores of 70-90 are expected to be well-modelled (good backbone), scores of 50-70 are low confidence, and scores of 0-50 should not be interpreted (likely disordered) ^70^. **(C)** To check for the presence of the NRPD1 clamp head domain, the cryo-EM density of the Arabidopsis Pol IV-RDR2 complex (EMD-31305) was inspected. The map was low-pass filtered to expose regions of lower resolution, which confirmed that the CYC-YPMF-containing domain is present. **(D)** The AlphaFold2 predicted NRPD1 clamp head domain is displayed according to its Coulombic electrostatic potential (ESP), with the color ranging from red for negative potential to blue for positive potential. The DNA facing part of the domain is largely positively charged, which is expected as DNA is largely negatively charged. For this analysis, the low confidence loop formed by residues Lys105 to Gln114 was trimmed because it was clashing with the DNA template.

**Figure S4.**
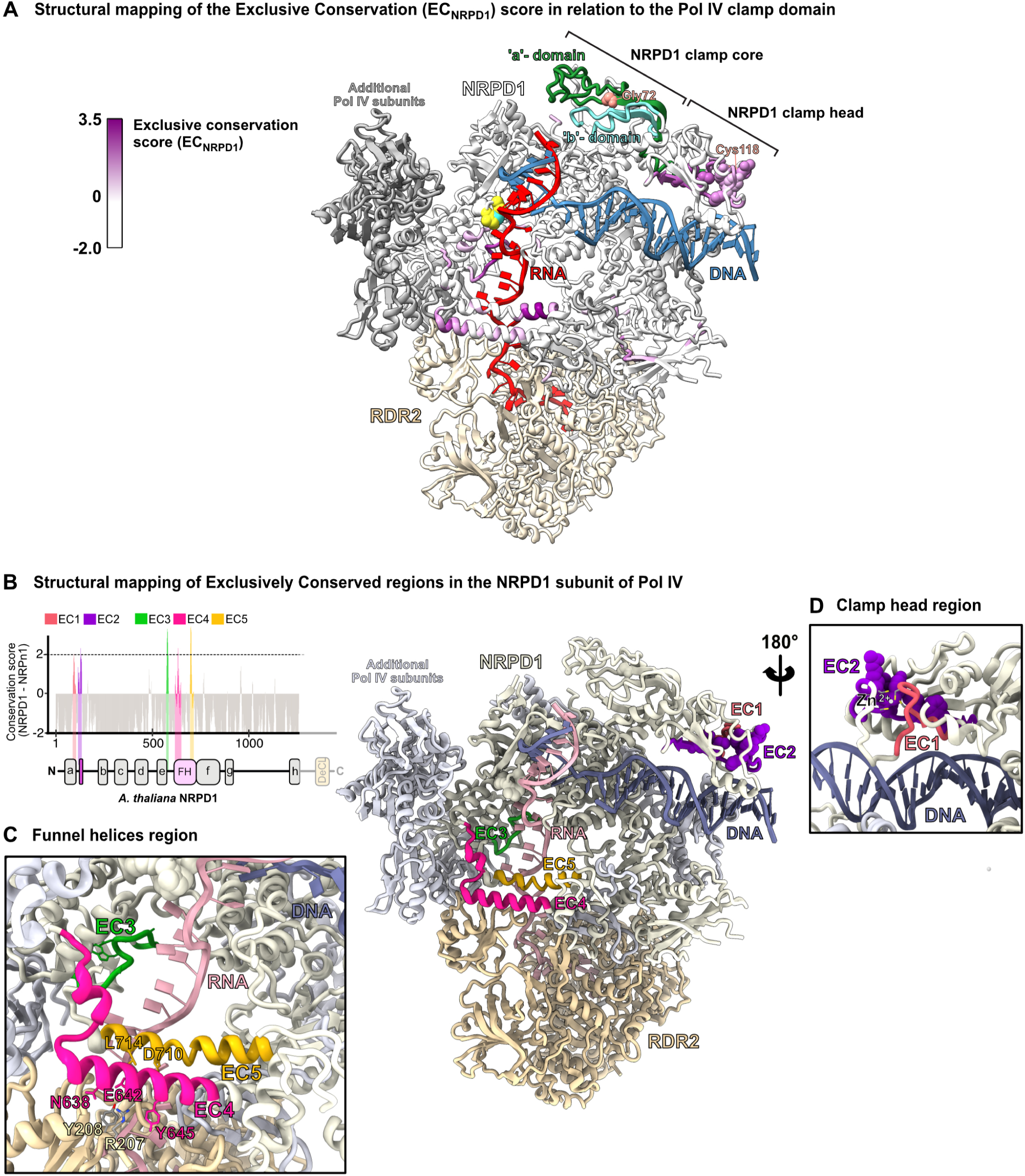
Structural representation of the exclusively conserved regions in the largest subunit of Pol IV (NRPD1). **(A)** Structural mapping of the NRPD1 exclusive conservation score (EC_NRPD1_, sliding average) rendered as a purple color gradient on the white NRPD1 amino acid chain. The DNA template is shown in blue, the transcript product of Pol IV in crimson, and the active site in yellow. Additional Pol IV subunits are shown in dark grey and RDR2 is shown in light beige. The universally conserved RNA polymerase ‘a’ and ‘b’-domains of NRPD1 are shown in green and turquoise, with the clamp core and clamp head domains indicated with brackets **(B)** The Pol IV-RDR2 model of Fig. 1E (right) but with the five peaks of NRPD1-exclusive conservation (EC1-EC5) plotted in color along the amino acid chain in the 3D model. Fig. 1D is reproduced at left as a key for comparing the universally conserved (‘a’-‘h’) domains to the funnel helices and EC regions. **(C)** Zoom inset of the NRPD1 funnel helices (EC4 and EC5) that interact with RDR2, as well as the EC3 loops that are adjacent to the Pol IV RNA transcript as it enters RDR2. Individual NRPD1 amino acids known to contact RDR2 (N638, E642, Y645, D710 and L714)^26^ are shown in bright pink and gold, indicating the identified EC4 and EC5 regions, respectively. Two amino acids from RDR2 that make contact with NRPD1 (R207 and Y208)^26^ are labelled in light beige. The NRPD1 amino acid positions covered by EC1-EC5 and relevant functional domains are provided in **Table S8**. **(D)** Zoom inset of the NRPD1 clamp head modeled using AlphaFold2, rotated 180 degrees with respect to the main model view and showing EC1 and EC2 regions, as well as Cys97-Cys100-Cys118-Cys121 coordinating zinc.

**Figure S5.**
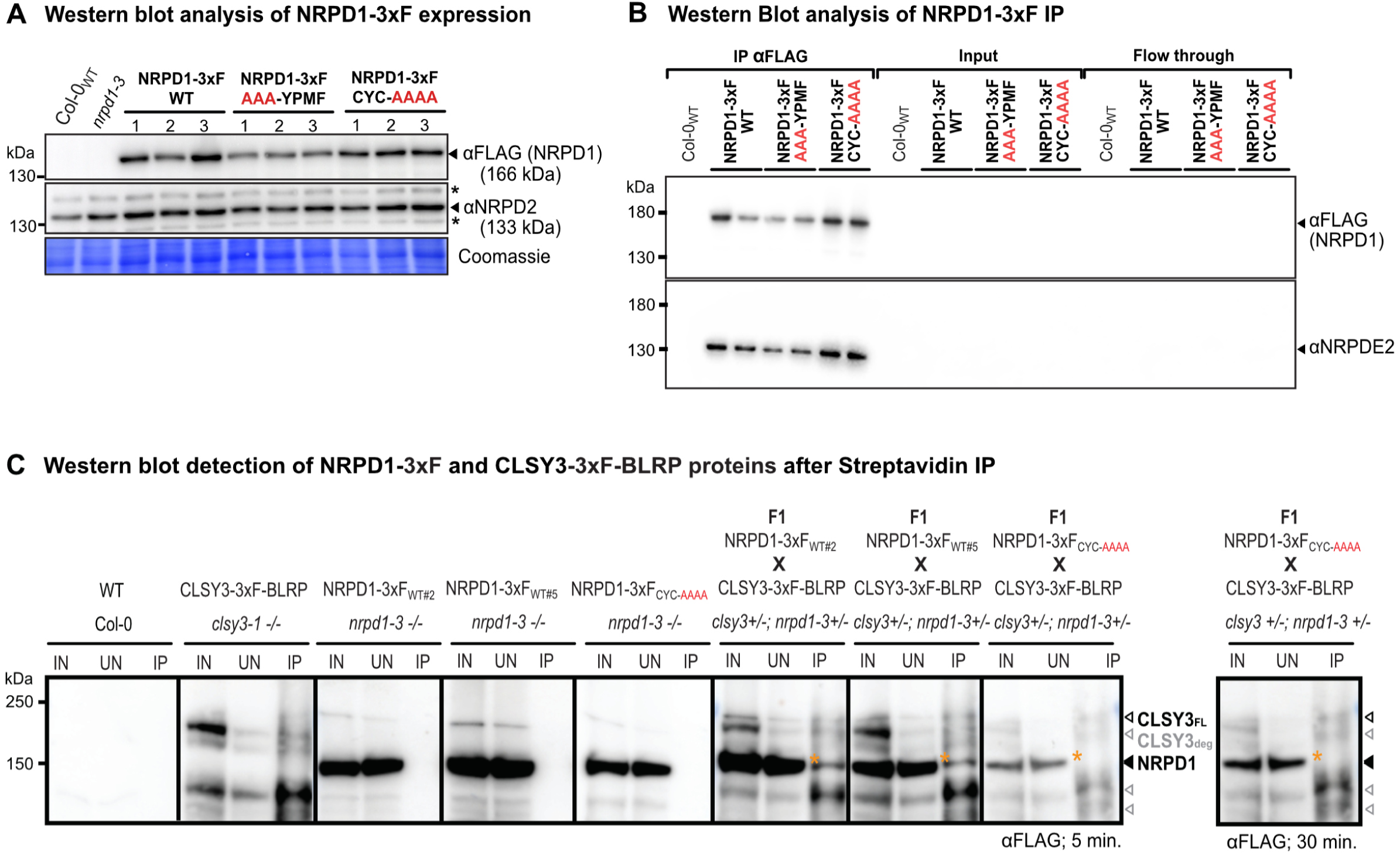
Western blot validation of Arabidopsis lines used for Pol IV complex IP-MS analyses. **(A)** Western blots of the NRPD1-3xF and NRPD2 subunits of Pol IV in extracts from Col-0_WT_ and *nrpd1-3* mutant flowers (negative controls) compared to independent insertion lines (1,2,3) expressing NRPD1-3xF_WT_, NRPD1-3xF_AAA-YPMF_ or NRPD1-3xF_CYC-AAAA_ variants in the *nrpd1-3* background (see Fig. 2A). **(B)** Western blots of the NRPD1-3xF variants and NRPD2 in the anti-FLAG immunopurification experiments analyzed by IP-MS (see Fig. 2B, C, D). The anti-FLAG immunopurified, input and column flow-through aliquots were loaded onto a single gel for comparison. (**C**) Anti-FLAG western blot detecting NRPD1 and CLSY3 proteins from either input (IN), unbound (UN), or streptavidin IP (IP) samples from the genotypes indicated above each blot. Protein sizes are indicated in kDa on the left and the bands corresponding to NPRD1 as well as both full length CLSY3 (CLSY3_FL_; Black) and several CLSY3 degradation products (CLSY3_deg_; Grey) are indicated on the right. The orange asterisk marks the location of the band representing the co-IP of NRPD1 with CLSY3. A longer exposure (30 min.) is shown to assess NRPD1 enrichment in the IP lane when similar levels of NRPD1 protein are present in the IN and UN lanes as observed for the NRPD1-3xF_WT#2_ and NRPD1-3xF_WT#5_ lines.

**Figure S6.**
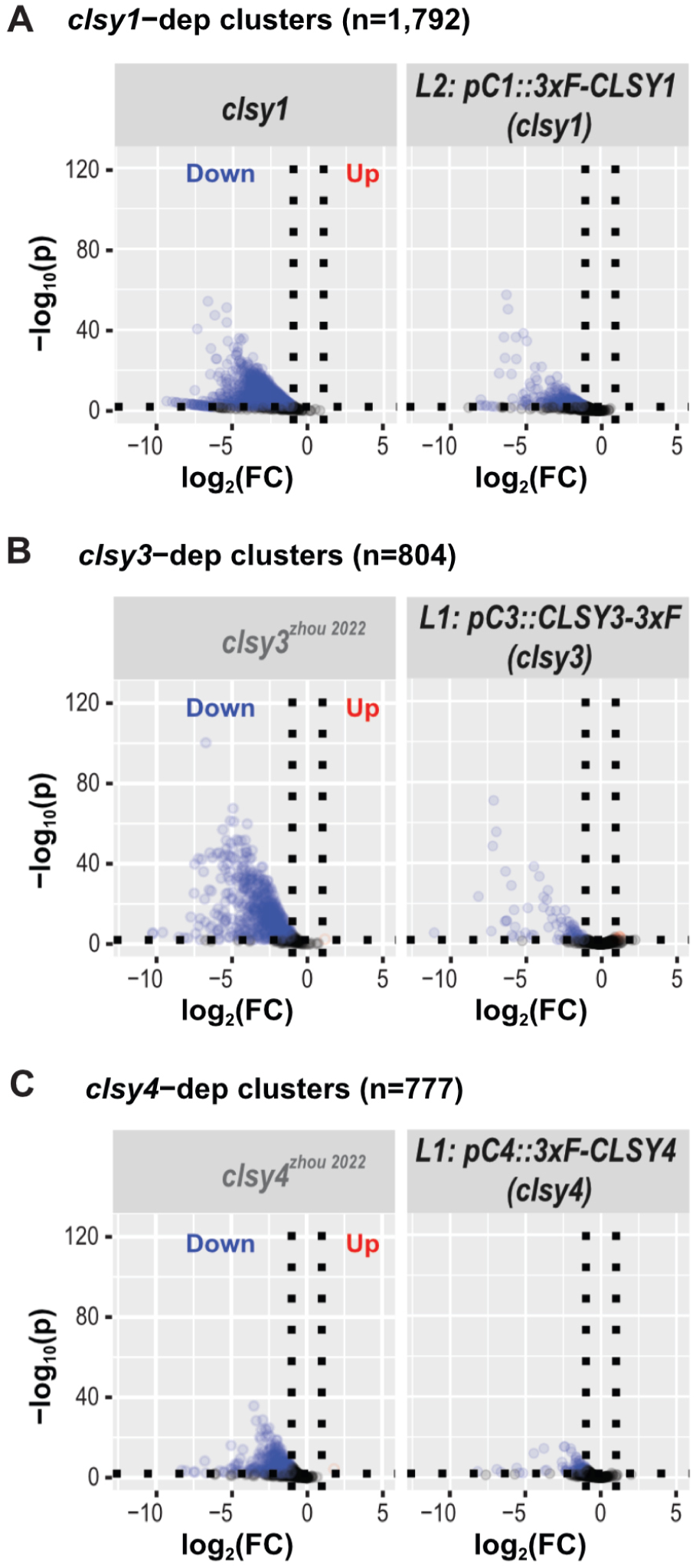
Complementation of *clsy* mutants with 3xF-tagged CLSY transgenes. (**A**-**C**) Volcano plots showing siRNA levels at *clsy1*-, *clsy3*-, and *clsy4*-dependent clusters (n=1792, 804, and 777, respectively), as defined in Zhou et al. 2018^10^. For each plot, clusters that are downregulated compared to wild-type controls (log_2_ FC ≤ −1 and FDR ≤ 0.01) are shown as blue circles, those unaffected are shown as black circles, and those upregulated (log_2_ FC ≥ 1 and FDR ≤ 0.01) are shown as red circles. For all panels, the plots with genotypes indicated in dark grey show previously published control data from Zhou et al. 2022^20^. The remaining plots show siRNA levels in the indicated *clsy* and *pol iv* mutants assayed in parallel with previously unpublished lines Zhou et al. 2022^20^. In each case, the *CLSY* genes are driven by their endogenous promoters (pC#) and the line number (L#) and N or C-terminal position of the 3xF tag is as indicated.

**Figure S7.**
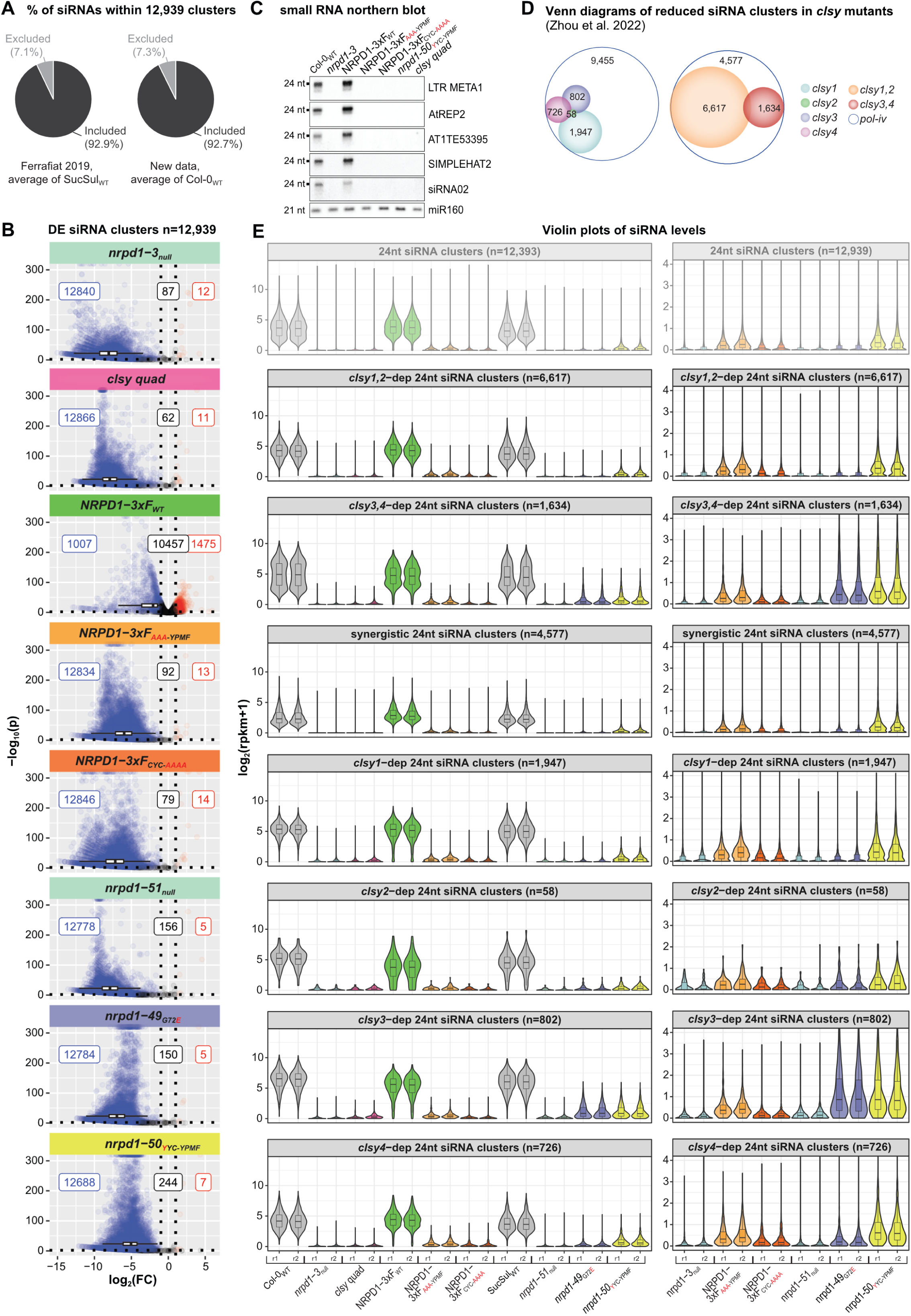
Assessment of siRNA levels in various *nrpd1* mutant and NRPD1-3xF variant lines. (**A**) Pie charts showing the fraction of 24 nt siRNAs from the indicated control samples that fall inside (included) or outside (excluded) of the 12,939 clusters defined in Zhou et al. 2022^20^. (**B**) Volcano plots showing 24 nt siRNA levels at the full set of clusters (n=12,939) defined in Zhou et al. 2022^20^. For each plot the genotype is indicated at the top and clusters that are downregulated compared to wild-type controls (log_2_FC ≤ −1 and FDR < 0.01) are shown as blue circles, those unaffected are shown as black circles, and those upregulated (log_2_ FC ≥ 1 and FDR < 0.01) are shown as red circles. The numbers of clusters in each category are indicated in the correspondingly colored boxes and a boxplot showing the overall effect of each mutant is included along the x-axis. (**C**) Northern blot analysis comparing small RNA levels in the indicated genotypes using probes for several RdDM targets (*LTR META1, AtREP2*, *AT1TE53395, SIMPLEHAT2* and *siRNA02*) or for miR160, as a loading control. (**D**) Scaled Venn diagrams depicting reduced siRNA clusters identified using flower tissue in the indicated mutants as originally published in Zhou et al. 2022^20^, except the numbers correspond to the total number of reduced siRNA clusters in each mutant rather than those unique to each mutant. (**E**) Violin plots showing the normalized expression levels [log_2_ (rpkm +1)] of 24nt siRNA clusters in the indicated categories and genotypes. Replicate samples are designated as r1 and r2. Select genotypes are shown on a smaller Y-axis scale on the right. The less opaque upper panel is reproduced from Fig. 4A.

### Supplementary tables

**Table S1.** List of the plant RNA polymerase subunits used for multiple sequence alignment and phylogenetic analyses (Fig. 1C, 1D, S1 and S2).

**Table S2.** Spectral counts of proteins identified by IP-MS of Pol IV complexes containing WT NRPD1-3xF_WT_, or the NRPD1-3xF_AAA-YPMF_ or NRPD1-3xF_CYC-AAAA_ variants.

**Table S3.** Spectral counts of proteins enriched in CLSY1, CLSY3 and CLSY4 3xF IP-MS experiments.

**Table S4.** Spectral counts of proteins copurifying with 3xHA-CLSY1 in NRPD1-3xF_WT_ or NRPD1-3xF_AAA-YPMF_ motif variant contexts identified by IP-MS.

**Table S5.** Summary of smRNA-seq data processing.

**Table S6.** Oligonucleotide primer and probe sequences.

**Table S7.** List of buffers used in this study.

**Table S8.** NRPD1 exclusively conserved regions and functional domains.

## References

1. Liu, P., Cuerda-Gil, D., Shahid, S. & Slotkin, R. K. The Epigenetic Control of the Transposable Element Life Cycle in Plant Genomes and Beyond. Annu. Rev. Genet. 56, 63–87 (2022).

2. Ozata, D. M., Gainetdinov, I., Zoch, A., O’Carroll, D. & Zamore, P. D. PIWI-interacting RNAs: small RNAs with big functions. Nat. Rev. Genet. 20, 89–108 (2019).

3. Zhou, M. & Law, J. A. RNA Pol IV and V in gene silencing: Rebel polymerases evolving away from Pol II’s rules. Curr. Opin. Plant Biol. 27, 154–164 (2015).

4. Rymen, B., Ferrafiat, L. & Blevins, T. Non-coding RNA polymerases that silence transposable elements and reprogram gene expression in plants. Transcription (2020) doi:10.1080/21541264.2020.1825906.

5. Haag, J. R. & Pikaard, C. S. Multisubunit RNA polymerases IV and V: purveyors of non-coding RNA for plant gene silencing. Nat. Rev. Mol. Cell Biol. 12, 483–492 (2011).

6. Law, J. A. et al. Polymerase IV occupancy at RNA-directed DNA methylation sites requires SHH1. Nature 498, 385–389 (2013).

7. Law, J. A. et al. A Protein Complex Required for Polymerase V Transcripts and RNA-Directed DNA Methylation in Arabidopsis. Curr. Biol. 20, 951–956 (2010).

8. Wongpalee, S. P. et al. CryoEM structures of Arabidopsis DDR complexes involved in RNA-directed DNA methylation. Nat. Commun. 10, 3916 (2019).

9. Law, J. A., Vashisht, A. A., Wohlschlegel, J. A. & Jacobsen, S. E. SHH1, a Homeodomain Protein Required for DNA Methylation, As Well As RDR2, RDM4, and Chromatin Remodeling Factors, Associate with RNA Polymerase IV. PLoS Genet. 7, e1002195 (2011).

10. Zhou, M., Palanca, A. M. S. & Law, J. A. Locus-specific control of the de novo DNA methylation pathway in Arabidopsis by the CLASSY family. Nat. Genet. 50, 865–873 (2018).

11. Haag, J. R. et al. In Vitro Transcription Activities of Pol IV, Pol V, and RDR2 Reveal Coupling of Pol IV and RDR2 for dsRNA Synthesis in Plant RNA Silencing. Mol. Cell 48, 811–818 (2012).

12. Ream, T. S. et al. Subunit Compositions of the RNA-Silencing Enzymes Pol IV and Pol V Reveal Their Origins as Specialized Forms of RNA Polymerase II. Mol. Cell 33, 192–203 (2009).

13. Andersen, P. R., Tirian, L., Vunjak, M. & Brennecke, J. A heterochromatin-dependent transcription machinery drives piRNA expression. Nature 549, 54–59 (2017).

14. Slotkin, R. K. et al. Epigenetic Reprogramming and Small RNA Silencing of Transposable Elements in Pollen. Cell 136, 461–472 (2009).

15. Ito, H. et al. An siRNA pathway prevents transgenerational retrotransposition in plants subjected to stress. Nature 472, 115–119 (2011).

16. Martins, L. M. & Law, J. A. Moving targets: Mechanisms regulating siRNA production and DNA methylation during plant development. Curr. Opin. Plant Biol. 75, 102435 (2023).

17. Smith, L. M. et al. An SNF2 Protein Associated with Nuclear RNA Silencing and the Spread of a Silencing Signal between Cells in Arabidopsis. Plant Cell 19, 1507–1521 (2007).

18. Knizewski, L., Ginalski, K. & Jerzmanowski, A. Snf2 proteins in plants: gene silencing and beyond. Trends Plant Sci. 13, 557–565 (2008).

19. Zhang, H. et al. DTF1 is a core component of RNA-directed DNA methylation and may assist in the recruitment of Pol IV. Proc. Natl. Acad. Sci. 110, 8290–8295 (2013).

20. Zhou, M. et al. The CLASSY family controls tissue-specific DNA methylation patterns in Arabidopsis. Nat. Commun. 13, 244 (2022).

21. Yang, D.-L. et al. Four putative SWI2/SNF2 chromatin remodelers have dual roles in regulating DNA methylation in Arabidopsis. Cell Discov. 4, 55 (2018).

22. Papareddy, R. K. et al. Chromatin regulates expression of small RNAs to help maintain transposon methylome homeostasis in Arabidopsis. Genome Biol. 21, 251 (2020).

23. Long, J. et al. Nurse cell--derived small RNAs define paternal epigenetic inheritance in Arabidopsis. Science 373, (2021).

24. Wang, Y. et al. ZMP recruits and excludes Pol IV-mediated DNA methylation in a site-specific manner. Sci. Adv. 8, eadc9454 (2022).

25. Singh, J., Mishra, V., Wang, F., Huang, H.-Y. & Pikaard, C. S. Reaction Mechanisms of Pol IV, RDR2, and DCL3 Drive RNA Channeling in the siRNA-Directed DNA Methylation Pathway. Mol. Cell 75, 576–589.e5 (2019).

26. Huang, K. et al. Pol IV and RDR2: A two-RNA-polymerase machine that produces double-stranded RNA. Science 374, 1579–1586 (2021).

27. Mishra, V. et al. Assembly of a dsRNA synthesizing complex: RNA-DEPENDENT RNA POLYMERASE 2 contacts the largest subunit of NUCLEAR RNA POLYMERASE IV. Proc. Natl. Acad. Sci. U. S. A. 118, (2021).

28. Blevins, T. et al. Identification of pol IV and RDR2-dependent precursors of 24 nt siRNAs guiding de novo DNA methylation in arabidopsis. Elife 4, 1–22 (2015).

29. Zhai, J. et al. A One Precursor One siRNA Model for Pol IV-Dependent siRNA Biogenesis. Cell 163, 445–455 (2015).

30. Wierzbicki, A. T., Ream, T. S., Haag, J. R. & Pikaard, C. S. RNA polymerase V transcription guides ARGONAUTE4 to chromatin. Nat. Genet. 41, 630–634 (2009).

31. Wierzbicki, A. T., Haag, J. R. & Pikaard, C. S. Noncoding Transcription by RNA Polymerase Pol IVb/Pol V Mediates Transcriptional Silencing of Overlapping and Adjacent Genes. Cell 135, 635– 648 (2008).

32. Xie, G. et al. Structure and mechanism of the plant RNA polymerase V. Science 379, 1209–1213 (2023).

33. Tucker, S. L., Reece, J., Ream, T. S. & Pikaard, C. S. Evolutionary history of plant multisubunit RNA polymerases IV and V: subunit origins via genome-wide and segmental gene duplications, retrotransposition, and lineage-specific subfunctionalization. Cold Spring Harb. Symp. Quant. Biol. 75, 285–297 (2010).

34. Trujillo, J. T., Seetharam, A. S., Hufford, M. B., Beilstein, M. A. & Mosher, R. A. Evidence for a Unique DNA-Dependent RNA Polymerase in Cereal Crops. Mol. Biol. Evol. 35, 2454–2462 (2018).

35. Luo, J. & Hall, B. D. A multistep process gave rise to RNA polymerase IV of land plants. J. Mol. Evol. (2007) doi:10.1007/s00239-006-0093-z.

36. Huang, Y. et al. Ancient Origin and Recent Innovations of RNA Polymerase IV and V. Mol. Biol. Evol. 32, 1788–1799 (2015).

37. Fukudome, A. et al. Structure and RNA template requirements of Arabidopsis RNA-DEPENDENT RNA POLYMERASE 2. Proc. Natl. Acad. Sci. U. S. A. 118, (2021).

38. Ferrafiat, L. et al. The NRPD1 N-terminus contains a Pol IV-specific motif that is critical for genome surveillance in Arabidopsis. Nucleic Acids Res. 47, 9037–9052 (2019).

39. Lahmy, S. et al. PolV(PolIVb) function in RNA-directed DNA methylation requires the conserved active site and an additional plant-specific subunit. Proc. Natl. Acad. Sci. U. S. A. 106, 941–946 (2009).

40. Haag, J. R., Pontes, O. & Pikaard, C. S. Metal A and metal B sites of nuclear RNA polymerases Pol IV and Pol V are required for siRNA-dependent DNA methylation and gene silencing. PLoS One 4, e4110 (2009).

41. Nudler, E. RNA polymerase active center: the molecular engine of transcription. Annu. Rev. Biochem. 78, 335–361 (2009).

42. Harlen, K. M. & Churchman, L. S. The code and beyond: transcription regulation by the RNA polymerase II carboxy-terminal domain. Nat. Rev. Mol. Cell Biol. 18, 263–273 (2017).

43. El-Shami, M. et al. Reiterated WG/GW motifs form functionally and evolutionarily conserved ARGONAUTE-binding platforms in RNAi-related components. Genes Dev. (2007) doi:10.1101/gad.451207.

44. Lahmy, S. et al. Evidence for ARGONAUTE4–DNA interactions in RNA-directed DNA methylation in plants. Genes Dev. 30, 2565–2570 (2016).

45. Coruh, C. et al. Comprehensive Annotation of Physcomitrella patens Small RNA Loci Reveals That the Heterochromatic Short Interfering RNA Pathway Is Largely Conserved in Land Plants. Plant Cell 27, 2148–2162 (2015).

46. Haag, J. R. et al. Functional Diversification of Maize RNA Polymerase IV and V Subtypes via Alternative Catalytic Subunits. Cell Rep. 9, 378–390 (2014).

47. Erhard, K. F. et al. RNA Polymerase IV Functions in Paramutation in Zea mays. Science 323, 1201–1205 (2009).

48. Xu, L. et al. Regulation of Rice Tillering by RNA-Directed DNA Methylation at Miniature Inverted-Repeat Transposable Elements. Mol. Plant (2020) doi:10.1016/j.molp.2020.02.009.

49. Ashkenazy, H. et al. ConSurf 2016: an improved methodology to estimate and visualize evolutionary conservation in macromolecules. Nucleic Acids Res. 44, (2016).

50. Cramer, P. et al. Architecture of RNA Polymerase II and Implications for the Transcription Mechanism. Science 288, 640–649 (2000).

51. Cramer, P., Bushnell, D. A. & Kornberg, R. D. Structural Basis of Transcription: RNA Polymerase II at 2.8 Angstrom Resolution. Science 292, 1863–1876 (2001).

52. Vos, S. M. et al. Structure of activated transcription complex Pol II-DSIF-PAF-SPT6. Nature 560, 607–612 (2018).

53. Schilbach, S., Aibara, S., Dienemann, C., Grabbe, F. & Cramer, P. Structure of RNA polymerase II pre-initiation complex at 2.9 Å defines initial DNA opening. Cell 184, 4064–4072.e28 (2021).

54. Ireland, S. M. & Martin, A. C. R. Zincbindpredict-Prediction of Zinc Binding Sites in Proteins. Molecules 26, (2021).

55. Hekkelman, M. L., de Vries, I., Joosten, R. P. & Perrakis, A. AlphaFill: enriching AlphaFold models with ligands and cofactors. Nat. Methods 20, 205–213 (2023).

56. Cavrak, V. V. et al. How a Retrotransposon Exploits the Plant’s Heat Stress Response for Its Activation. PLoS Genet. 10, e1004115 (2014).

57. Thieme, M. et al. Inhibition of RNA polymerase II allows controlled mobilisation of retrotransposons for plant breeding. Genome Biol. 18, 134 (2017).

58. Antosz, W. et al. The Composition of the Arabidopsis RNA Polymerase II Transcript Elongation Complex Reveals the Interplay between Elongation and mRNA Processing Factors. Plant Cell 29, 854–870 (2017).

59. Ream, T. S. et al. Subunit compositions of Arabidopsis RNA polymerases I and III reveal Pol I- and Pol III-specific forms of the AC40 subunit and alternative forms of the C53 subunit. Nucleic Acids Res. 43, 4163–4178 (2015).

60. Yariv, B. et al. Using evolutionary data to make sense of macromolecules with a ‘face-lifted’ ConSurf. Protein Sci. 32, e4582 (2023).

61. Zhang, H.-W. et al. A cryo-EM structure of KTF1-bound polymerase V transcription elongation complex. Nat. Commun. 14, 3118 (2023).

62. Chow, H. T. & Mosher, R. A. Small RNA-mediated DNA methylation during plant reproduction. Plant Cell 35, 1787–1800 (2023).

63. Shahzad, Z., Eaglesfield, R., Carr, C. & Amtmann, A. Cryptic variation in RNA-directed DNA-methylation controls lateral root development when auxin signalling is perturbed. Nat. Commun. 11, 218 (2020).

64. Borges, F. et al. Loss of Small-RNA-Directed DNA Methylation in the Plant Cell Cycle Promotes Germline Reprogramming and Somaclonal Variation. Curr. Biol. 31, 591–600.e4 (2021).

65. Onodera, Y. et al. Plant Nuclear RNA Polymerase IV Mediates siRNA and DNA Methylation-Dependent Heterochromatin Formation. Cell 120, 613–622 (2005).

66. Herr, A. J., Jensen, M. B., Dalmay, T. & Baulcombe, D. C. RNA Polymerase IV Directs Silencing of Endogenous DNA. Science 308, 118–120 (2005).

67. Greenberg, M. V et al. Identification of genes required for de novo DNA methylation in Arabidopsis. Epigenetics 6, 344–354 (2011).

68. Link, D. et al. Functional characterization of the Beet necrotic yellow vein virus RNA-5-encoded p26 protein: evidence for structural pathogenicity determinants. J. Gen. Virol. 86, 2115–2125 (2005).

69. Goddard, T. D. et al. UCSF ChimeraX: Meeting modern challenges in visualization and analysis. Protein Sci. 27, 14–25 (2018).

70. Jumper, J. et al. Highly accurate protein structure prediction with AlphaFold. Nat. 2021 5967873 596, 583–589 (2021).

71. Meng, E. C., Pettersen, E. F., Couch, G. S., Huang, C. C. & Ferrin, T. E. Tools for integrated sequence-structure analysis with UCSF Chimera. BMC Bioinformatics 7, 339 (2006).

72. Faurobert, M., Pelpoir, E. & Chaïb, J. Phenol Extraction of Proteins for Proteomic Studies of Recalcitrant Plant Tissues. Methods Mol. Biol. 355, 9–14 (2007).

73. Hughes, C. S. et al. Single-pot, solid-phase-enhanced sample preparation for proteomics experiments. Nat. Protoc. 14, 68–85 (2019).

74. Jami-Alahmadi, Y., Pandey, V., Mayank, A. K. & Wohlschlegel, J. A. A Robust Method for Packing High Resolution C18 RP-nano-HPLC Columns. J. Vis. Exp. (2021) doi:10.3791/62380.

75. Wang, Q. et al. The blue light-dependent phosphorylation of the CCE domain determines the photosensitivity of Arabidopsis CRY2. Mol. Plant 8, 631–643 (2015).

76. Zhang, Y. et al. The Use of Variable Q1 Isolation Windows Improves Selectivity in LC-SWATH-MS Acquisition. J. Proteome Res. 14, 4359–4371 (2015).

77. Cox, J. & Mann, M. MaxQuant enables high peptide identification rates, individualized p.p.b.-range mass accuracies and proteome-wide protein quantification. Nat. Biotechnol. 26, 1367–1372 (2008).

78. Tsou, C.-C. et al. DIA-Umpire: comprehensive computational framework for data-independent acquisition proteomics. Nat. Methods 12, 258–64, 7 p following 264 (2015).

79. Kong, A. T., Leprevost, F. V, Avtonomov, D. M., Mellacheruvu, D. & Nesvizhskii, A. I. MSFragger: ultrafast and comprehensive peptide identification in mass spectrometry-based proteomics. Nat. Methods 14, 513–520 (2017).

80. Wickham, H. ggplot2: Elegant Graphics for Data Analysis. (Springer-Verlag, 2016).

81. Kassambara, A. ggpubr v0.6.0: ‘ggplot2’ Based Publication Ready Plots. R package. (2023).

82. Böhrer, M. et al. Integrated Genome-Scale Analysis and Northern Blot Detection of Retrotransposon siRNAs Across Plant Species. in Methods in Molecular Biology 387–411 (2020). doi:10.1007/978-1-0716-0712-1_23.

83. Martin, M. Cutadapt Removes Adapter Sequences From High-Throughput Sequencing Reads. EMBnet.journal 17, (2011).

84. Axtell, M. J. ShortStack: comprehensive annotation and quantification of small RNA genes. RNA 19, 740–751 (2013).

85. Barnett, D. W., Garrison, E. K., Quinlan, A. R., Strömberg, M. P. & Marth, G. T. BamTools: a C++ API and toolkit for analyzing and managing BAM files. Bioinformatics 27, 1691–1692 (2011).

86. Heinz, S. et al. Simple combinations of lineage-determining transcription factors prime cis-regulatory elements required for macrophage and B cell identities. Mol. Cell 38, 576–589 (2010).

87. Li, H. et al. The Sequence Alignment/Map format and SAMtools. Bioinformatics 25, 2078–2079 (2009).

88. Love, M. I., Huber, W. & Anders, S. Moderated estimation of fold change and dispersion for RNA-seq data with DESeq2. Genome Biol. 15, 550 (2014).

